# Genome-Wide Large-Scale Multi-Trait Analysis Characterizes Global Patterns of Pleiotropy and Unique Trait-Specific Variants

**DOI:** 10.1101/2022.06.03.494686

**Authors:** Guanghao Qi, Surya B. Chhetri, Debashree Ray, Diptavo Dutta, Alexis Battle, Samsiddhi Bhattacharjee, Nilanjan Chatterjee

## Abstract

Genome-wide association studies (GWAS) have found widespread evidence of pleiotropy, but characterization of global patterns of pleiotropy remain highly incomplete due to insufficient power of current approaches. We develop fastASSET, an extension of the method ASSET, to allow computationally efficient detection of variant-level pleiotropic association across a large number of traits. We analyze GWAS summary statistics of 116 complex traits of diverse types collected from the NIH GRASP repository and a number of other large GWAS consortia. We identify a total of 2,293 independent loci at the genome-wide significance level and found that the lead variants in nearly all of these loci (∼99%) to be associated with to two or more (median = 6) traits. Further, the estimated degree of pleiotropy for the detected variants strongly predicted their degree of pleiotropy across a much larger number of traits (K=4,114) in the UK Biobank Study. Follow-up analyses of 21 unique trait-specific variants suggest that they are often linked to the expression in trait-related tissues for a small number of genes, some of which are well known to be involved in relevant biological processes. Our findings provide deeper insight into the nature of complex trait pleiotropy and leads to, for the first time, identification of highly unique trait-specific susceptibility variants.

## Introduction

Genome-wide association studies (GWAS) have identified thousands of susceptibility loci across individual complex traits and diseases^1^. Studies have also pointed to the evidence of widespread pleiotropy^2–5^, i.e., genetic variants within individual loci are often associated with multiple traits. The discovery of pleiotropy has transformed the analysis and interpretation of GWAS data. It has, for example, led to the development of more powerful statistical methods for association testing^6–11^ and polygenic prediction^10^, robust methods for causal inference accounting for pleiotropic associations^12–15^, as well as new study designs to investigate multiple or even hundreds of traits simultaneously^16–19^. The presence of pleiotropy also poses unique opportunities and challenges for developing or/and repurposing of drugs while minimizing their “off-target” antagonistic effects^20^.

There has been long a quest in genetics to characterize degree and nature of pleiotropy for complex traits^21^. Early population genetic models postulated universal pleiotropy where genetic variants at any given locus has the potential to affect all traits^21^. More recent experimental studies, however, have suggested that while pleiotropy is highly prevalent, it is likely to be modular in nature, i.e. any given gene is likely to affect relatively a small number of traits^21^. Data from recent GWAS for human traits have also indicated pleiotropy is pervasive^2–5^, but quantifying the true extent of pleiotropy has been challenging. A number of recent studies^4,22^ have quantified degree of pleiotropy associated with a variant based on the number of associated traits that reach genome-wide significance (*p* < 5 × 10 ^−8^) in individual trait analysis, but such analysis inevitably lead to serious underestimation of the extent of pleiotropy due to lack of power for GWAS of individual traits for the detection of smaller effect-sizes. Another recent study adopted use of a more liberal threshold (z-statistic > 2)^23^ for large scale detection of pleiotropy, but such an analysis can introduce a large number of false positives.

In addition, while previous studies have mostly focused on detecting highly pleiotropic loci and variants^17,22,24^, we believe that given the evidence of highly abundant pleiotropy, an interesting line of investigation would be to detect highly trait-specific loci and variants and explore their unique biological characteristics, if any. Identification of trait specific genetic association may facilitate identification of “core genes” under the omnigenic model for complex traits^25,26^, distinguish the genetic architecture of related traits and potentially help develop drug targets with less side effects. Detection of trait-specific associations, however, requires highly powerful methods for detecting pleiotropy as undetected weaker associations would lead to increase in findings for trait-specific variants. The most comprehensive analysis of pleiotropy based on current GWAS^4^ has reported that almost 70% of identified individual SNPs to be associated with a single trait-domain, but these are likely to be highly overestimated due to the lack of power of the underlying analytic method.

In this paper, we develop fastASSET, an extension of the ASsociation analysis based on subSETs (ASSET), which allows detection of any association between a variant and an underlying subset of traits that contribute to the association signal ^7^. A major advantage of ASSET, compared to other multi-trait association tests^6,8,9^, is that it not only allows powerful detection of SNPs that show any association across a group of traits, but also readily maps significant SNPs to sets of associated traits. The subset selection feature, which has shown to have robust sensitivity for the detection of true traits under association even when power of individual studies vary^7^, makes the method ideally suited for the investigation of the extent of pleiotropy in current GWAS. While the method has been successfully applied to a number of multi-trait GWAS analyses^27–29^ involving a limited number of traits, it is not feasible to implement for the analysis of very large number of traits because of computational burdens associated with all subset search. Here, we develop fastASSET that allows association testing for individual SNPs across a large number of traits by first incorporating a pre-screening step and then performing ASSET on selected traits with suitable adjustment for the pre-screening step.

We use fastASSET to analyze 116 traits collected from large GWAS Consortia and the Genome-Wide Repository of Associations Between SNPs and Phenotypes (GRASP) hosted by the National Institutes of Health (NIH). We identify 2,293 independent loci that are associated with at least one trait and show that lead variants at nearly all of these loci are associated with two or more traits. We show that the degree of pleiotropy we estimate for the underlying variants based on the 116 traits predicts strongly the level of their pleiotropy associated with a much larger number of traits (>4,000) in the UK Biobank Study. We conduct a series of follow-up analysis to examine whether the degree of pleiotropy of genetic variants may be related to functional mechanisms, including cis-regulatory effects, chromatin states, transcription factor (TF) binding, and enhancer-gene connections. We further provide detail characterization of 21 highly unique trait-specific variants, i.e., those were associated with only one trait in the fastASSET analysis.

## Results

### Overview of datasets and methods

We collect 338 summary-level datasets from the NIH GRASP repository, and supplement it with 20 summary-level datasets from large GWAS consortia. See **Figure 1** for data preprocessing pipeline and **Methods** for details. After filtering out duplicated and highly correlated traits, removing studies with small sample size and data quality issues, we retain 116 well-powered studies consisting of primarily participants of European ancestry (**Supplementary Tables 1 and 2**). The studies cover a wide range of complex traits and diseases in 16 domains (**Supplementary Table 1**). Genetic correlation analysis using linkage disequilibrium (LD) score regression reveals widespread genome-wide pleiotropy (**Supplementary Figure 1**). We restrict further analysis to 7,462,466 single-nucleotide polymorphisms (SNP) for which summary-statistics were available for at least 50 out of the 116 traits and have minor allele frequency (MAF) > 0.01.

**Figure 1.**
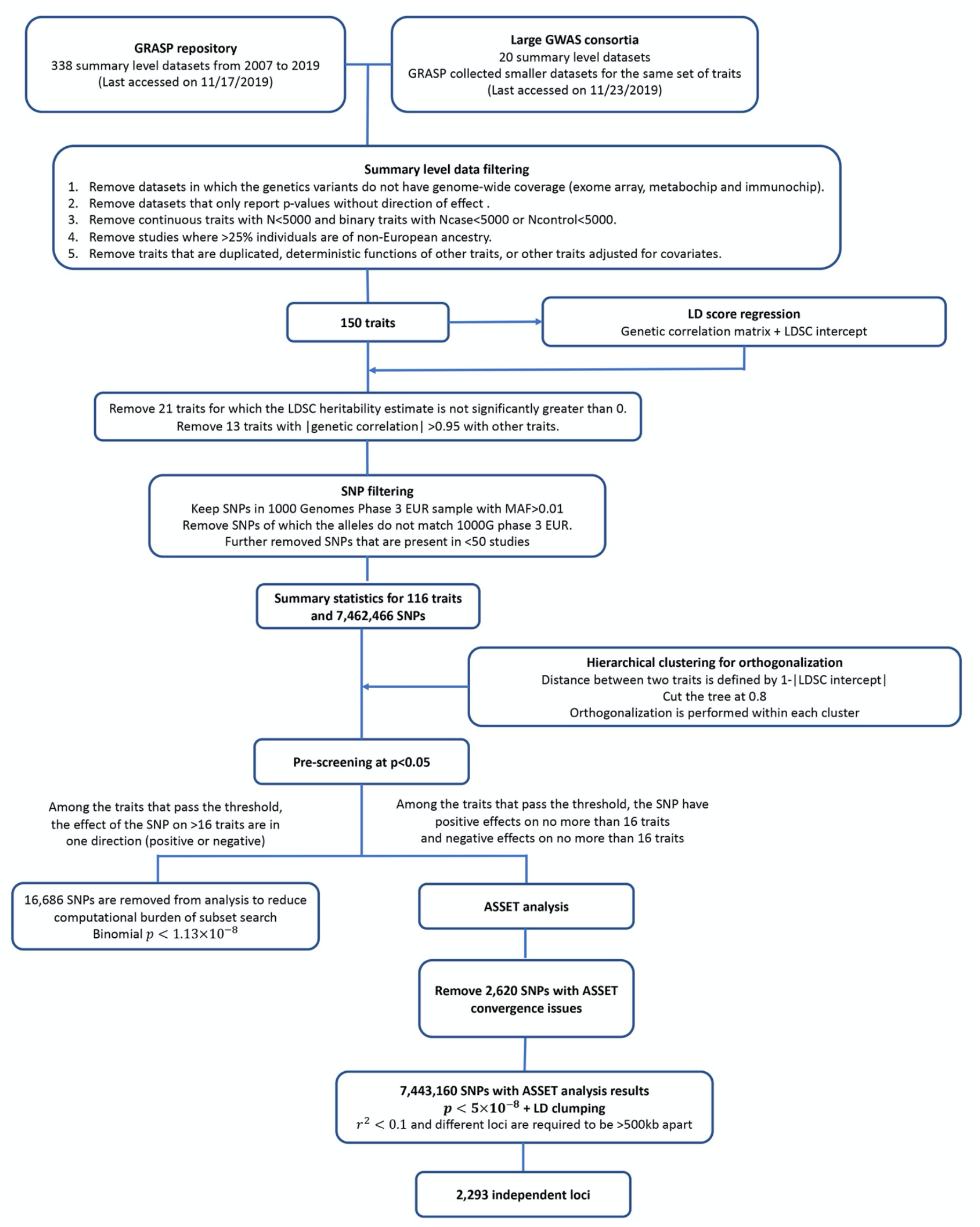
Data preprocessing and analysis pipeline. Details are provided in Methods.

We develop fastASSET, an extension of the ASSET method^7^, to conduct multi-trait association testing across a large number of traits (see **Methods and Supplementary Notes** for details). ASSET was originally designed to conduct single-SNP association tests by performing meta-analysis across all subsets of traits and then evaluating the significance of the maximum of meta-analysis z-scores over all subsets^7^. In this method, for SNPs which reach a desired significance threshold for association, the set of traits for which the underlying meta-analysis z-statistics is maximized defines the set of underlying associated traits. For the analysis of a large number of traits, the original ASSET, which searches through all subsets, is computationally intractable. The fastASSET method reduces the computational burden by only searching among the traits which show suggestive evidence of associations. The method first de-correlates the z-statistics for a given SNP associated with different traits using estimates of phenotypic correlations available from the LD score regression (see **Online Methods**). Next, it selects the set of traits that shows suggestive level of association (e.g. *p* < 0.05) with the given SNP based on the de-correlated z-statistics. The de-correlated z-statistics are then adjusted for the pre-selection step as 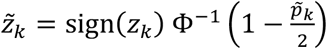 where 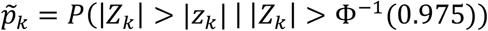 and Φ is the quantile function of standard normal distribution. The adjusted z-statistics are then further transformed back to the original scale of the traits, also using estimates of phenotypic correlations available from the LD score regression, and then these Z-statistics are incorporated as input into the original ASSET method. For each SNP, fastASSET outputs a p-value for the association with any trait under consideration (“global association”) and a set of associated traits. Simulation studies show the method has well calibrated type-I error (**Supplementary Table 3**) and robust power for detection of global association when the sample size is large (e.g. *N* ≥ 50*k*, **Supplementary Figure 2**). Since our dataset has a median sample size of >100k, fastASSET is likely to be powerful for our application. Across most of the scenarios, fastASSET estimates the level of pleiotropy more accurately than previous methods based on the number of associated traits that reach *p* < 5 × 10^−8^ or *p* < 0.05 in individual trait analysis (**Supplementary Figure 3**). It tends to outperform the other methods when the fraction of true non-null traits is small (2 or 5 out of 116 traits), but can lead to underestimation when this number is large (e.g. 15 out of 116 traits, **Supplementary Figure 3**). See **Supplementary Notes** for description of the simulation settings. The method has been incorporated into the ASSET R package (see Code Availability for more details).

### Quantifying and validating levels of pleiotropy

We find widespread genetic associations and varying degree of pleiotropy across the genome (**Figure 2 and Supplementary Figure 4**). Identified associations represent 2,293 loci (fastASSET p-value < 5 × 10^−8^, *r*^2^ < 0.1 and >500kb apart, **Supplementary Table 4**). We further investigate the signal density within each locus by counting the number of independent genome-wide significant SNPs (fastASSET p-value < 5 × 10^−8^, *r*^2^ < 0.1) within 100kb of the lead SNP (see **Methods** for details). We observe substantial variation in signal density (**Figure 2**). Multiple independent signals were present at nearly half (48% out of 2,293) of the loci; 11% of the loci harbored at least 5 independent signals and some loci can harbor up to 25 signals (**Figure 2**). For the loci with multiple associated SNPs, levels of pleiotropy can vary within the locus (**Figure 2**). For example, SNP rs3760047 (chr16:281299) is associated with 4 traits, but 8 out of the 15 significant SNPs within 100kb are associated with 2-5 traits and the remaining 7 SNPs are associated with 6-10 traits (**Supplementary Table 4**). In the following, we use the lead SNPs detected by fastASSET for each locus to study the level of pleiotropy across the genome and its relationship with different types of variant annotations.

**Figure 2.**
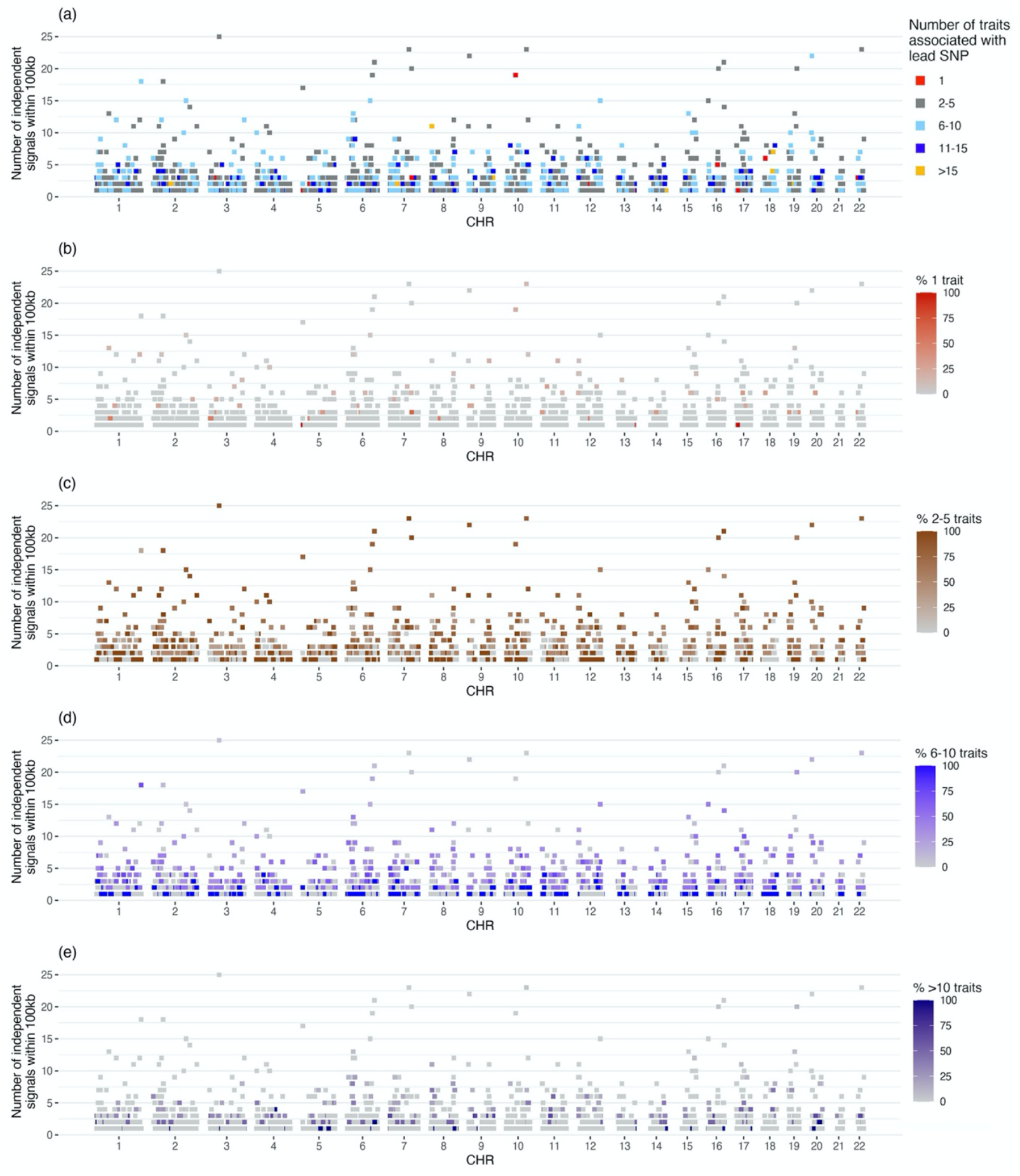
Brisbane plot showing signal density and levels of pleiotropy of 2,293 genome-wide significant loci identified by fastASSET analysis across 116 traits. We define significant SNPs as those that have fastASSET p-value < 5 × 10^−8^ and clump them to obtain 10,628 independent SNPs (*r*^2^ < 0.1), which we then group into 2,293 loci whose lead SNPs (strongest association within a locus) are at least 500kb apart. Each dot represents the lead SNP of a locus, and the y axis is the number of independent significant SNPs within 100kb from the lead SNP. The dots are colored by the number of traits associated with the lead SNP as reported by fastASSET (a) or the proportion of significant SNPs in a locus (within 100kb from the lead SNP) that fall into each category of pleiotropy: trait-specific (b – associated with 1 trait), low pleiotropy (c – associated with 2-5 traits), medium pleiotropy (d – associated with 6-10 traits) or high pleiotropy (e – associated with >10 traits). Nearly half (1,094) of the loci harbor at least two independent genome-wide significant SNPs within 100kb of the lead SNP (including the lead SNP itself); 248 (11%) loci harbor at least 5 significant SNPs within 100kb of the lead SNP.

The vast majority of the lead SNPs are associated with 2-10 traits with a median of 6 (**Figure 3a**). At the two ends of the spectrum, we found 21 SNPs to be associated with only one trait, representing highly trait-specific genetic mechanisms and 58 SNPs to be highly pleiotropic defined as those which are associated with 16 or more traits. Next, we investigate whether the degree of pleiotropy we estimate for the lead variants based on the 116 traits also predicts degree of pleiotropy that the same variants will manifest across a much wider spectrum of traits. To test this hypothesis, we collect the summary statistics for 4,114 traits from the Neale lab UK Biobank (UKB) GWAS^30,31^, and quantify the levels of pleiotropy for each of the 2,293 lead SNPs by the number of associated traits at per-SNP false discovery rate (FDR) < 0.05. We observe that degree of pleiotropy estimated from our study strongly predicts that observed in the UK Biobank (spearman correlation=0.43, p-value < 2.2 × 10^− 16^, **Figure 3b**). The relationship remains highly significant even after adjusting for LD score (partial correlation=0.42, p-value < 2.2 × 10^−16^). The analysis indicates that pleiotropic characteristics of the detected SNPs is not specific to the selected traits in our discovery analysis and likely represent a much broader property related to their roles in gene regulation.

**Figure 3.**
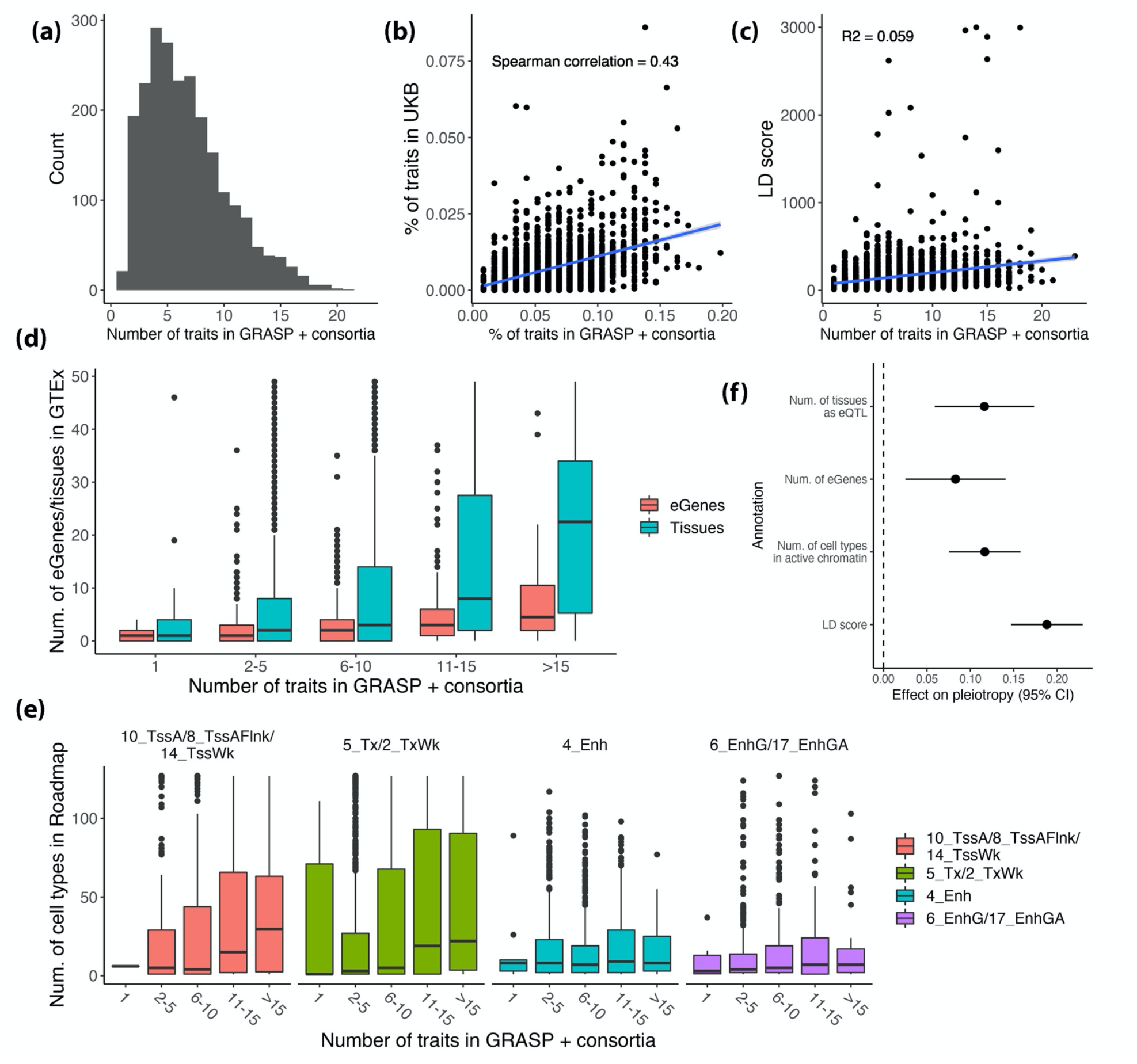
Number of associated traits detected by fastASSET and its relationship with different variant annotations. Results are shown for only the lead SNPs of 2,293 independent loci identified by fastASSET. Sub-figures show (a) Distribution of number of associated traits; (b) Relationship with the number of associated traits in the UK Biobank (per-SNP FDR < 0.05, each point represents a lead SNP); (c) Relationship with linkage disequilibrium (LD score); (d) Relationship with number of tissues and genes for which a SNP is a significant eQTL in GTEx v8; (e) Relationship with the number of cell types in which the SNP is in each active chromatin state; (f) Effect of each annotation on pleiotropy (number of associated traits) conditional on other annotations. Chromatin states are learned by IDEAS method (Zhang et al, 2017) using data from the Roadmap Epigenomics Consortium. Coefficients in (f) are estimated using multiple linear regression model of the form *pleiotropy* ∼(*LD score)* + (*num. of tissues as eQTL)* + (*num. of eGenes)* + (*num. of cell types in active chromatin*) where all dependent and independent variables are standardized to have unit variance.

### Relationship with functional and LD annotations

We investigated how the level of pleiotropy is correlated with different types of variant annotations. First, we found that the numbers of associated traits detected by fastASSET to be positively correlated with LD score values across the 2,293 lead SNPs (R^2^=0.059, p-value = 4.95 × 10^−32^, **Figure 3c**). The pattern is expected since lead SNPs which tag more SNPs due to LD will appear to be associated with a larger number of traits when there are distinct causal variants for distinct traits within the loci. We also found that SNPs associated with a larger number of traits tend to be significant expression quantitative trait loci (eQTL) for a larger number tissues and for a larger number of eGenes (both R^2^=0.069, p-value < 2.2 × 10^−16^). Trait-specific SNPs are significant eQTL for a median of 1 tissue and 1 gene, while highly pleiotropic SNPs (>15 traits) are significant eQTL for a median of 22.5 tissues and 4.5 genes (**Figure 3d**). While such pattern has been reported earlier^4^, we observe a much sharper dose-response trend and higher level of statistical significance (e.g. compared to Figure 1e in reference ^4^), arising likely due to the higher accuracy of the fastASSET analysis for the detection of degree of pleiotropy. We also found more pleiotropic SNPs tend to be in regions of active chromatin state in a larger number of tissue or cell types, reflected by results from the IDEAS method^32,33^ (**Figure 3e**). This trend is especially pronounced for the promoter (10_TssA, 8_TssAFlnk and 14_TssWk) and transcription (5_Tx and 2_TxWk) related chromatin states, but weaker for enhancer-related states (4_Enh, 6_EnhG and 17_EnhGA). We observe similar relationship using chromatin states learned by ChromHMM (**Supplementary Figure 5**). In a multivariate regression analysis that accounts for LD and all functional annotations simultaneously, we found the relationship between degree of pleiotropy and three types of functional characteristics of the SNPs each remain highly significant (p-value < 0.005), with eQTL tissue specificity showing the largest effect size in per standard deviation unit (**Figure 3f**). Further, analysis based on the JASPAR and HOCOMOCO databases^34,35^ revealed that more pleiotropic SNPs (>10 traits) are more likely to overlap with transcription factor binding sites (TFBS) (p-value=0.022, **Supplementary Figure 6**). The relationship remains significant (p-value=0.021) after adjusting for the annotations in Figure 3 (**Supplementary Figure 6**). Finally, analysis based on activity-by-contact (ABC) model revealed that among the SNPs that overlap with enhancers, level of pleiotropy increases with the number of active tissues (correlation=0.09, p-value=0.069, **Supplementary Figure 6)**. However, this relationship disappears after adjusting for other annotations (p-value=0.65, **Supplementary Figure 6**).

### Trait-specific Variants

The fastASSET analysis identified 21 independent trait-specific lead SNPs, defined as those for which associated subset included only one trait (**Table 1**). We observe that the association p-value between a trait-specific SNP and the primary trait are at least 10^8^ times lower than the next most strongly associated trait (**Figure 4 and Supplementary Figure 7**). Highly pleiotropic SNPs, however, often have comparable level of associations with a large number of traits (**Figure 4 and Supplementary Figure 7**). We also validate the nature of trait specificity of these SNPs in the external UK Biobank study (**Table 1**). Notably, even though UK Biobank covered a much larger of traits, the trait-specific SNPs we identified for Alzheimer’s disease, breast cancer, prostate cancer, Crohn’s disease (CD) are only associated with traits related to these diseases, except rs4631223 detected for primary trait CD was also found to be associated with Urea. Some of the disease-specific SNPs we detected were not significant in the UK Biobank (**Table 1**) likely due to its lower power compared to large case-control studies that contributed to our discovery analysis. For quantitative traits, we observe replication of trait-specificity in the UK biobank for most of the trait-specific SNPs for intelligence, male baldness, age at menarche and monocyte count (**Table 1**). The trait-specific SNPs we detected for heel bone mineral density (BMD), height and diastolic blood pressure (DBP), however, were associated with a larger number of traits in the UKB, but the majority of the additional traits were related to the primary trait.

**Table 1.**
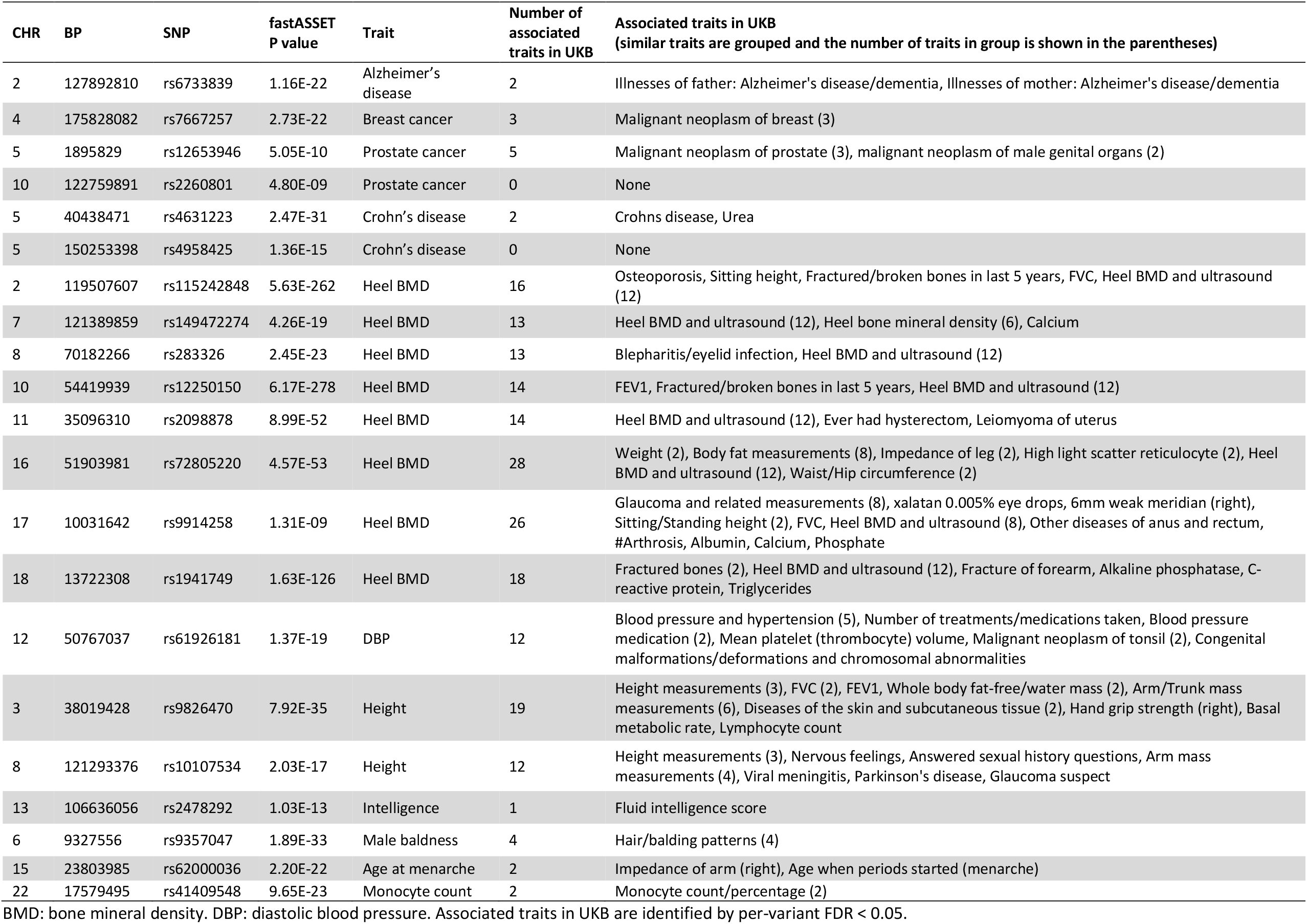
Association of trait-specific lead SNPs identified by fastASSET analysis with UK Biobank Traits.

| CHR | BP | SNP | fastASSET<br>P value | Trait | Number of<br>associated<br>traits in UKB | Associated traits in UKB<br>(similar traits are grouped and the number of traits in group is shown in the parentheses) |
| --- | --- | --- | --- | --- | --- | --- |
| 2 | 127892810 | rs6733839 | 1.16E-22 | Alzheimer's disease | 2 | Illnesses of father: Alzheimer's disease/dementia, Illnesses of mother: Alzheimer's disease/dementia |
| 4 | 175828082 | rs7667257 | 2.73E-22 | Breast cancer | 3 | Malignant neoplasm of breast (3) |
| 5 | 1895829 | rs12653946 | 5.05E-10 | Prostate cancer | 5 | Malignant neoplasm of prostate (3), malignant neoplasm of male genital organs (2) |
| 10 | 122759891 | rs2260801 | 4.80E-09 | Prostate cancer | 0 | None |
| 5 | 40438471 | rs4631223 | 2.47E-31 | Crohn's disease | 2 | Crohns disease, Urea |
| 5 | 150253398 | rs4958425 | 1.36E-15 | Crohn's disease | 0 | None |
| 2 | 119507607 | rs115242848 | 5.63E-262 | Heel BMD | 16 | Osteoporosis, Sitting height, Fractured/broken bones in last 5 years, FVC, Heel BMD and ultrasound (12) |
| 7 | 121389859 | rs149472274 | 4.26E-19 | Heel BMD | 13 | Heel BMD and ultrasound (12), Heel bone mineral density (6), Calcium |
| 8 | 70182266 | rs283326 | 2.45E-23 | Heel BMD | 13 | Blepharitis/eyelid infection, Heel BMD and ultrasound (12) |
| 10 | 54419939 | rs12250150 | 6.17E-278 | Heel BMD | 14 | FEV1, Fractured/broken bones in last 5 years, Heel BMD and ultrasound (12) |
| 11 | 35096310 | rs2098878 | 8.99E-52 | Heel BMD | 14 | Heel BMD and ultrasound (12), Ever had hysterectomy, Leiomyoma of uterus |
| 16 | 51903981 | rs72805220 | 4.57E-53 | Heel BMD | 28 | Weight (2), Body fat measurements (8), Impedance of leg (2), High light scatter reticulocyte (2), Heel BMD and ultrasound (12), Waist/Hip circumference (2) |
| 17 | 10031642 | rs9914258 | 1.31E-09 | Heel BMD | 26 | Glaucoma and related measurements (8), xalatan 0.005% eye drops, 6mm weak meridian (right), Sitting/Standing height (2), FVC, Heel BMD and ultrasound (8), Other diseases of anus and rectum, #Arthrosis, Albumin, Calcium, Phosphate |
| 18 | 13722308 | rs1941749 | 1.63E-126 | Heel BMD | 18 | Fractured bones (2), Heel BMD and ultrasound (12), Fracture of forearm, Alkaline phosphatase, C-reactive protein, Triglycerides |
| 12 | 50767037 | rs61926181 | 1.37E-19 | DBP | 12 | Blood pressure and hypertension (5), Number of treatments/medications taken, Blood pressure medication (2), Mean platelet (thrombocyte) volume, Malignant neoplasm of tonsil (2), Congenital malformations/deformations and chromosomal abnormalities |
| 3 | 38019428 | rs9826470 | 7.92E-35 | Height | 19 | Height measurements (3), FVC (2), FEV1, Whole body fat-free/water mass (2), Arm/Trunk mass measurements (6), Diseases of the skin and subcutaneous tissue (2), Hand grip strength (right), Basal metabolic rate, Lymphocyte count |
| 8 | 121293376 | rs10107534 | 2.03E-17 | Height | 12 | Height measurements (3), Nervous feelings, Answered sexual history questions, Arm mass measurements (4), Viral meningitis, Parkinson's disease, Glaucoma suspect |
| 13 | 106636056 | rs2478292 | 1.03E-13 | Intelligence | 1 | Fluid intelligence score |
| 6 | 9327556 | rs9357047 | 1.89E-33 | Male baldness | 4 | Hair/balding patterns (4) |
| 15 | 23803985 | rs62000036 | 2.20E-22 | Age at menarche | 2 | Impedance of arm (right), Age when periods started (menarche) |
| 22 | 17579495 | rs41409548 | 9.65E-23 | Monocyte count | 2 | Monocyte count/percentage (2) |

BMD: bone mineral density. DBP: diastolic blood pressure. Associated traits in UKB are identified by per-variant FDR < 0.05.

**Figure 4.**
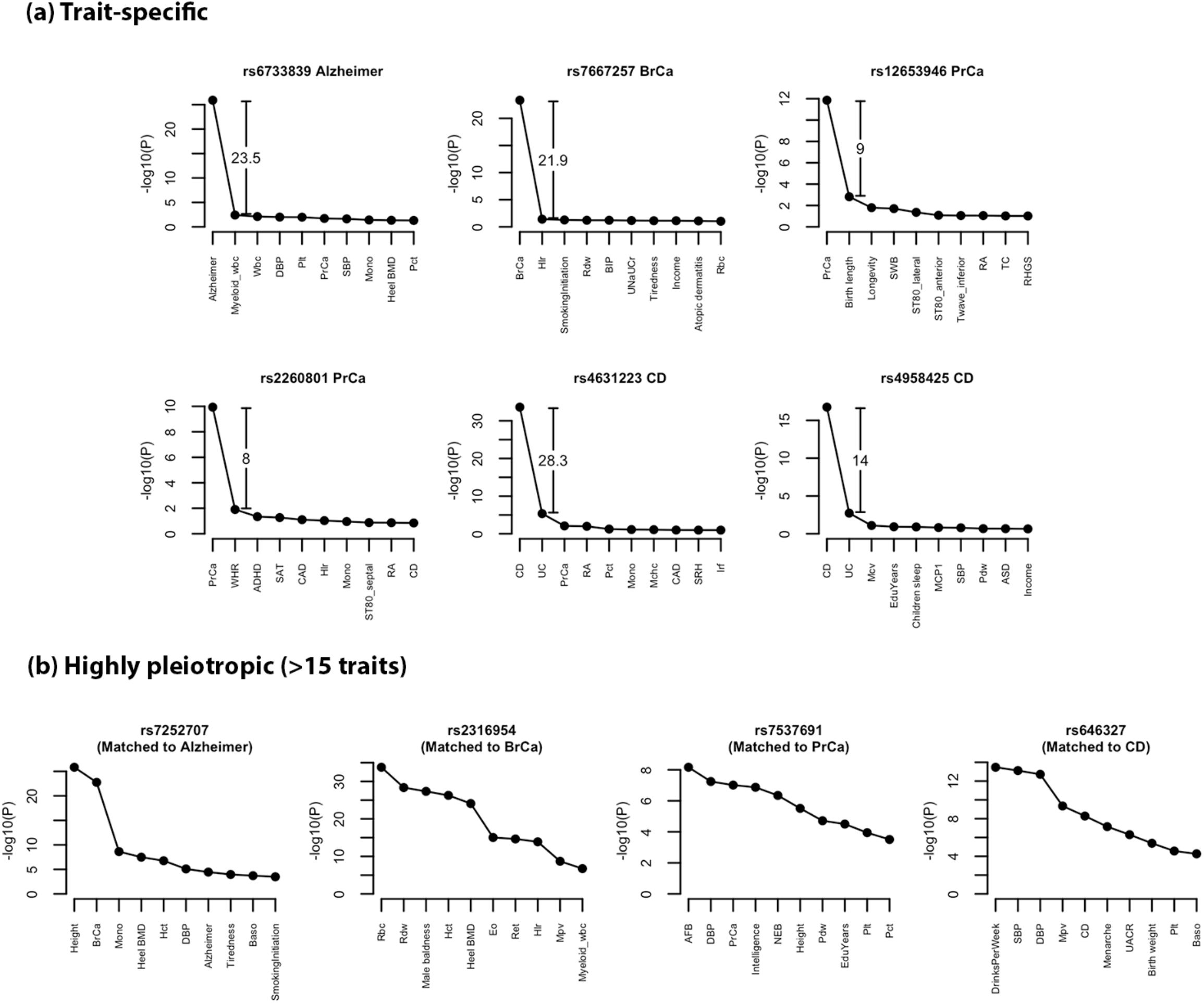
Top 10 associated traits for trait-specific SNPs for binary diseases and matched highly pleiotropic SNPs. Traits are ordered by descending order of -log10(p-value) in each panel. We show the trait-specific SNPs for binary diseases as examples. Each disease-specific SNP is matched to the highly pleiotropic SNP (>15 traits) that has the smallest association p-value with the disease (matched trait shown in the parentheses). See Supplementary Figures 7 for trait-specific SNPs for quantitative traits and matched highly pleiotropic SNPs. See **Methods** for details of the matching procedure.

### Cis-regulatory effects for trait-specific SNPs

To gain further insight into the mechanisms driving trait-specific SNPs, we search for their *cis*-eGenes and corresponding tissues in GTEx^36^, as well as the biological functions of the eGenes in the GeneCards database^37^. Among the 21 trait-specific SNPs, 11 SNPs are eQTLs for at least one gene-tissue pair (q-value < 0.05) in GTEx v8 (**Figure 5 and Supplementary Figure 8**). Five of them are associated with the expression of one single gene. rs6733839 (Alzheimer’s disease) is a significant eQTL for *BIN1* gene in aorta (**Figure 5**). Though aorta does not appear to be the relevant tissue for Alzheimer’s disease, the SNP is also moderately associated the expression of the *BIN1* in brain cerebellum (**Supplementary Figure 9**). rs7667257 (breast cancer) is an eQTL for only *GLRA3* in breast mammary tissue, which is exactly the relevant issue for breast cancer. rs12653946 (prostate cancer) is associated with the expression of *IRX4* in 4 tissues but has the largest effect size in prostate – the disease’s tissue of occurrence (**Figure 5**). rs9914258 (heel BMD) is associated with the expression of *GAS72* with the strongest effect in cultured fibroblasts (**Supplementary Figure 8**). rs41409548 (monocyte count) is associated with the expression of *IL17RA* in brain cortex (**Supplementary Figure 8**) and a number of other brain tissues (**Supplementary Figure 10**).

**Figure 5.**
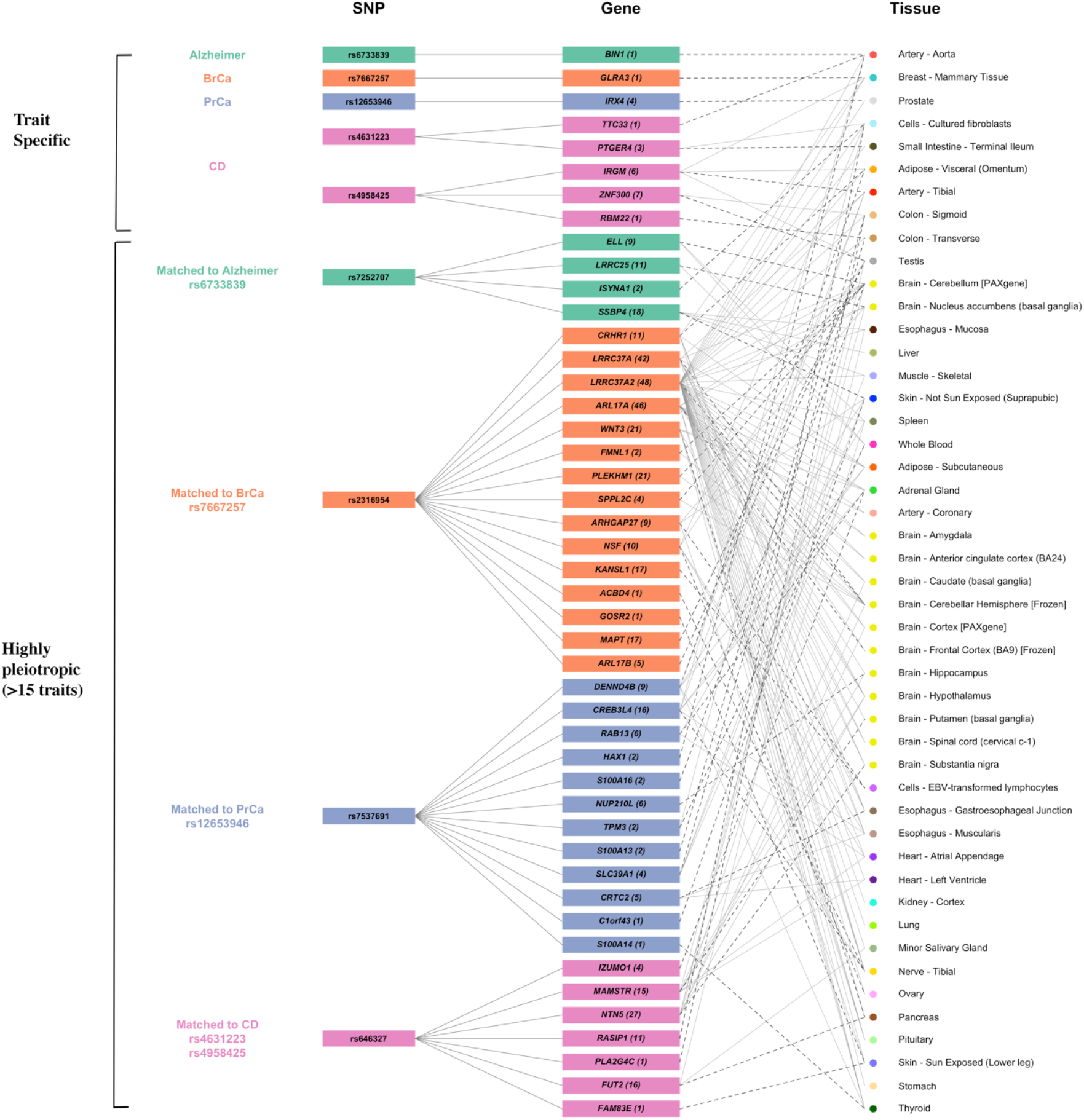
Cis-regulatory effects of trait-specific SNPs for binary diseases and matched highly pleiotropic SNPs. For each SNP, we show its protein-coding eGenes (q-value < 0.05) and the corresponding top tissues in GTEx v8. The top tissues for each variant-gene pair are defined as those with eQTL effect size is >0.7*(the largest effect size among significant tissues for this variant-gene pair); the tissue harboring the largest effect is highlighted by darker dashed lines. In addition, we annotate each gene name by the total number of associated tissues (q-value < 0.05) regardless of effect size. Each trait-specific SNP is matched to the highly pleiotropic SNP (>15 traits) that have the strongest association with the trait (lowest p-value). See **Methods** for details of the matching procedure. Pseudogenes and non-coding RNAs are excluded. See Supplementary Figure 8 for trait-specific SNPs for quantitative traits and highly pleiotropic SNPs. BrCa: breast cancer; PrCa: prostate cancer; CD: Crohn’s disease.

The other trait-specific SNPs, though do not show highly specific gene or tissue specific eQTLs, are generally associated with a small number of genes in a small number of tissues compared to highly pleiotropic “matched” SNPs that are also associated with the same traits. rs4631223 (Crohn’s disease) is associated with the expression of *TTC33* and *PTGER4*, and the largest effect size for *PTGER4* is in small intestine (**Figure 5**), the organ where Crohn’s disease usually occurs. Another trait-specific SNP for Crohn’s disease, rs4958425, is associated with the expression of *IRGM, ZNF300* and *RBM22*. The association for *RMB22* occurs in colon transverse and a secondary association for *ZNF300* occurs in colon sigmoid (**Figure 5**), both of which are in the digestive system. Though the strongest associations for *IRGM* do not occur in digestive tissues, it is moderately associated with rs4958425 in small intestine (**Supplementary Figure 11**). Previous studies have supported the connections between the trait mechanisms and the functions of some of the genes above, including the role of BIN1 in Alzheimer’s disease, *PTGER4* and *IRGM* in Crohn’s disease and *IL17RA* in monocyte count ^38–41^ (**Supplementary Table 5**).

We further used COLOC^42^ to conduct colocalization analysis for the loci indexed by 11 trait-specific SNPs that are also significant eQTLs in GTEx. We find evidence of colocalization (PP4>0.8) between genetic effects on the trait and on gene expression for 8 trait-specific SNPs (**Supplementary Table 6**), indicating shared causal SNPs. Many of the genes and tissues we highlighted as potential mechanisms for trait-specific SNPs (**Figure 5 and Supplementary Figure 8**) are also supported by colocalization, e.g. *IRX4* and prostate, *PTGER4* and small intestine, *RBM22* and transverse colon, etc. (**Supplementary Table 6**). These results provide further evidence that trait-specificity may be driven by gene- and tissue-specific cis-regulatory effects.

### Chromatin state of trait-specific SNPs

We explore another potential mechanism of trait-specificity using chromatin states learned by the IDEAS method^32^ using data from Roadmap Epigenomics Consortium^43^. With a few exceptions, we find that trait specific SNPs tend to be in active chromatin state only in specific tissue or cell types (**Figure 6**). On the other hand, highly pleiotropic SNPs tend to have active chromatin state in a wide range of tissue and cell types (**Figure 6**). For a number of trait-specific SNPs, chromatin state results strongly link the SNP to trait/disease related tissues. For example, rs4631223 (Crohn’s disease) is in active chromatin state in blood and T-cells, B-cells and digestive tissues, which are related to the immune mechanism of Crohn’s disease and the organ where it occurs. rs6733839 (Alzheimer’s disease) is active in brain and muscle tissues (**Figure 6**), both of which were shown to be involved the disease^44–46^, though it is also active in immune cells and digestive tissues, whose role in the disease is less clear. rs61926181 (DBP) is active in heart tissues which is also closely related to DBP.

**Figure 6.**
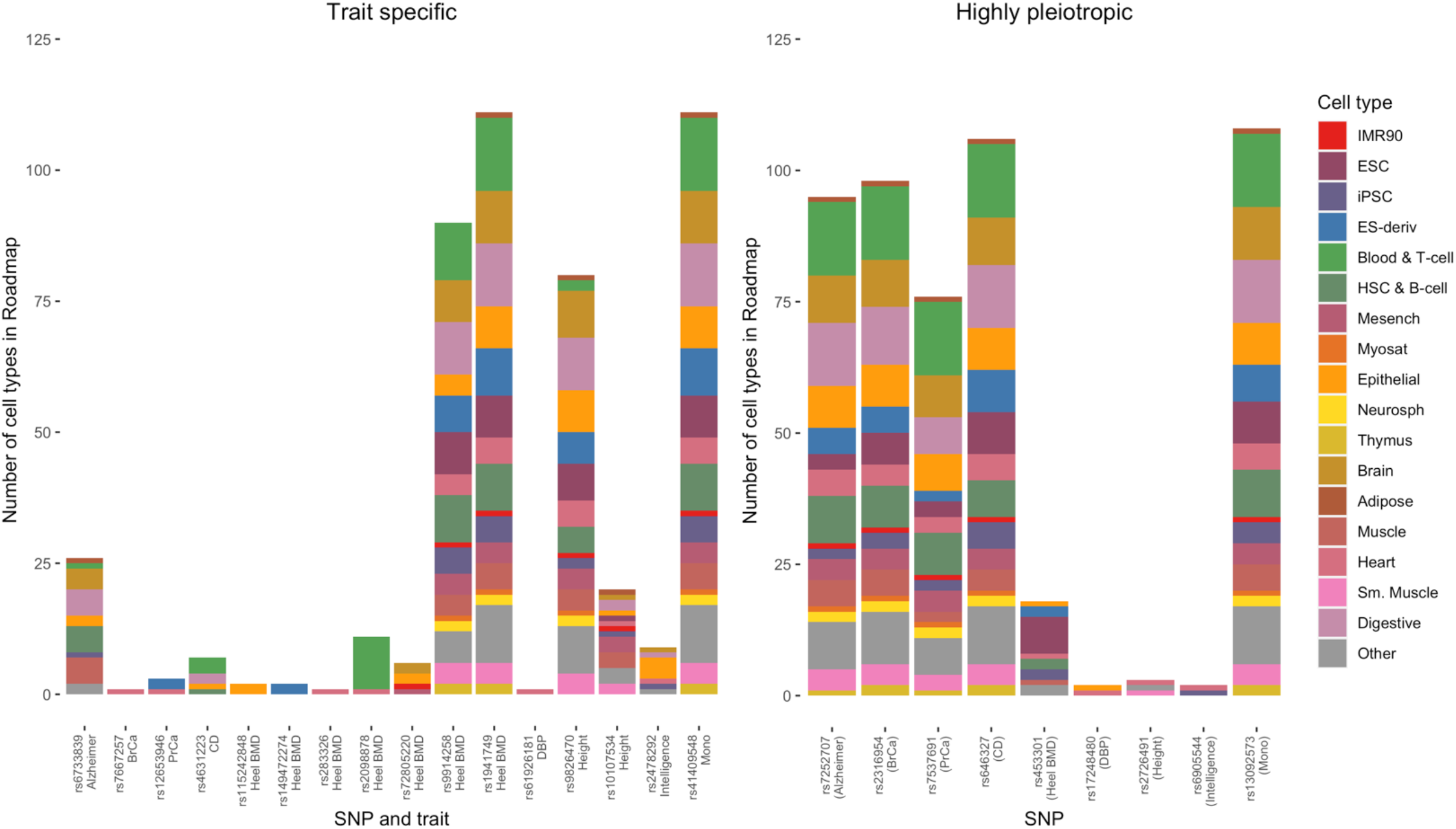
Cell types in which the trait-specific and matched highly pleiotropic SNPs have active chromatin state. Each trait-specific SNP is matched to the highly pleiotropic SNP (>15 traits) that have the strongest association with the trait (lowest p-value). See **Methods** for details of the matching procedure. Chromatin states are learned by applying the IDEAS method to data from the Roadmap Epigenomics Project. BrCa: breast cancer; PrCa: prostate cancer; CD: Crohn’s disease; BMD: bone mineral density; DBP: diastolic blood pressure; mono: monocyte count.

### Trait specificity and effect sizes

We explored whether effect-size of variants on lead traits have any relationship with the degree of their pleiotropy. Under the omnigenic model^25,26^ for complex traits, a small number of “core genes” have direct effects on the trait, and a large number of “peripheral genes” affect the trait by regulating the core genes. Under this model, SNPs near the core genes of a trait could be expected to show the largest effect sizes and more trait-specific effects. On the contrary, if the SNPs with the largest effect sizes for a given trait are also highly pleiotropic, then they could be near master regulators of many pathways and the core genes, if existent, could be indistinguishable from peripheral genes. We investigate the hypothesis for three types of traits: 1) complex binary diseases 2) blood and urine biomarkers 3) anthropometric and social behavioral traits (**Supplementary Figure 12**). For each trait, we select a list of top genome-wide significant SNPs (single-trait p-value < 5 × 10^−8^, see **Supplementary Figure 12** for details) and explore the relationship between rank of their standardized regression coefficients and the number of additional traits these variants are associated with. While in general we did not see a strong relationship, for a number of traits, including schizophrenia, body mass index (BMI), height, waist-hip-ratio (WHR), neuroticism and lipid levels, the SNPs with larger effect sizes on the given trait were more pleiotropic than SNPs with smaller effect-sizes (**Supplementary Figure 12**).

## Discussion

In this paper, we develop fastASSET and use it to conduct a large-scale multi-trait genome-wide association analysis across 116 traits. We quantify degree of pleiotropy at the level of individual SNPs across the genome and their relationship with various annotation characteristics. Specifically, we detect a much higher degree of pleiotropy at individual variant level than earlier studies and show that patterns of pleiotropy may be driven by multiple functional mechanisms. In a first of its kind effort, we identify 21 highly trait-specific variants and conduct extensive follow-up studies to show that they can often be linked to the functions of specific genes in trait-related tissues.

While our analysis confirms the ubiquitous nature of pleiotropy reported in prior studies^4,22^, we are able to provide several novel insights due to both the use of a rapid powerful method for cross-trait association analysis and detailed follow-up investigation of relationship between the level of pleiotropy and functional characteristics across the SNPs. First, we observed that pleiotropy is extremely ubiquitous, not only at a locus level but also at the level of individual variants. A prior large-scale study of pleiotropy across 558 traits based on UK Biobank had noted earlier that while >90% of loci are pleiotropic, about 60% of the individual SNPs show pleiotropic association with more than one trait and only 32% show association across multiple domains of traits^4^. Our analysis, though based on a smaller number of traits, reveals that lead variants across almost all the identified loci are associated with more than one trait. The nature of pleiotropy, however, is “modular” is nature^21^, i.e. a given variant has detectable effect only on a small fraction of the traits studied.

Second, our study confirms observations reported in previous studies^4,36^ that variants affecting expression in multiple genes and tissues tend to be more pleiotropic, but we are able to detect the trend at a much stronger resolution due to more precise characterization of the nature of pleiotropy. We further show that the level of pleiotropy is associated with the tissue-specificity of active chromatin states, and the relationship appears to be stronger for promoter and transcription related states than enhancers. Among SNPs located in enhancer regions, we found pleiotropy can be driven by interaction with multiple genes in multiple cell types. Finally, we also demonstrate enrichment of TF binding sites among pleiotropic SNPs.

Finally, a unique contribution of this paper is the identification of the highly unique trait-specific variants. We show that the trait-specific SNPs detected by fastASSET have dominant associations with the primary trait that are far stronger than any secondary associations (**Figure 4 and Supplementary Figure 7**) and they show similar pattern of trait specificity even when examined against a much larger number of traits in the UK Biobank study (**Table 1**). Despite their small number, the trait-specific SNPs can provide unique insight to biological mechanisms. We find that they often have clearly interpretable regulatory effects (**Figures 5 and 6**). Some of them regulate the expression of a single gene in a single tissue. Others are active in multiple tissues, but the strongest regulatory effects occur in trait-related tissues. For some other SNPs the mechanisms are less obvious, but still clearly contrast with highly pleiotropic SNPs which can affect many genes in many tissues.

Motivated by the omnigenic model for complex traits, we also explore whether SNPs that have larger effect sizes for a given trait, and possibly near core genes, are more likely to show trait specific effect compared to SNPs that are associated with the same trait but weaker effect-sizes (**Supplementary Figure 12**). For some traits, including cancer and autoimmune diseases, we observe suggestive but weak evidence of such a pattern. On the contrary, for several other traits including psychiatric diseases, anthropometric and social science traits, we observed a fairly distinct but opposite trend in that the variants with the largest effects tend to be more pleiotropic. Thus it appears that for complex traits, especially those which show highest degree of polygenicity, pattens of common variant associations may not fit well the hypothesis that SNPs near trait specific core genes will show the strongest effect sizes.

Our study has several limitations. In our discovery analysis, we restricted the analysis to a diverse but relatively small set of (K=116) traits that have large associated GWAS (N ranging from 7,000 to 1.2 million; >50% studies have N>100k). As we have excluded vast number of other traits for which GWAS data are also available, we have not been able to provide a more complete genotype-phenotype maps. Nevertheless, we observe that the level of pleiotropy we observe for the detected SNPs with respect to a smaller number of traits strongly correlates with the level of pleiotropy observed in the UK Biobank study with respect to a much larger number of traits. It is particularly notable that the trait-specific SNPs we detect based on only 116 traits largely show same or similar trait specific effect when validated against more than 4,000 traits in the UK Biobank.

Another limitation of our study is that we have primarily identified pleiotropic associations based on single SNP association analysis without accounting for LD within loci. In single-SNP association analysis, variant-level pleiotropy may arise due to LD even when there are distinct causal variants associated with distinct traits within a genomic locus. In our follow-up analysis we show that while estimated degree of pleiotropy is indeed correlated with LD score across the variants, its relationship with various functional characteristics of the genome could not be explained by LD alone. A more detailed characterization of pleiotropy at the level of each individual variant will ultimately require joint fine-mapping analysis across many traits by each locus, but methods for large-scale multi-trait fine-mapping analysis are currently not available.

In summary, we use a powerful new method to carry out a large-scale pleiotropic analysis across GWAS of a diverse set of traits. Our study provides novel insights into functional characteristics of the genome that contribute to pleiotropy and leads to the identification of unique trait-specific genetic variants which have not been previously explored. In the future, large-scale cross-trait fine-mapping studies are needed to pinpoint causal variants and the underlying nature of pleiotropy.

## Methods

### Data and initial filtering

We collect 338 full summary level datasets published between 2007 and 2019 from the NIH Genome-Wide Repository of Associations Between SNPs and Phenotypes (GRASP). GRASP includes a wide range of phenotypes including anthropometric traits, biomarkers, blood cell levels, adipose volume, early growth traits, social science Indices, cardiometabolic diseases, psychiatric and neurological diseases, autoimmune diseases, etc. We further collect 20 summary-level datasets from GWAS consortia for the traits included in GRASP but when the consortia offer a study with larger sample sizes. See **Supplementary Tables 1 and 2** for a full list of datasets. Thus restricting the analysis to the GRASP traits allowed us to carry out the pleiotropic analysis of across a diverse set of traits with large available GWAS, but avoid including many overlapping traits such as those in UK Biobank.

We apply several filtering steps sequentially (see **Figure 1** for a flow chart): 1) remove datasets in which the genetics variants do not have genome-wide coverage (genotyped by exome array, metabochip and immunochip); 2) remove datasets that only report p-values without direction of effect; 3) remove studies with small sample size: for continuous traits, we remove studies with sample size < 5,000; for binary traits, we remove studies with < 5,000 cases or < 5,000 controls; 4) remove studies where >25% individuals are of non-European ancestry; 5) remove duplicated traits: among the studies for the same trait, we keep the study with the largest sample size and discard the rest of them; 6) remove traits that are deterministic functions of other traits in our study, or other traits adjusted of covariates. After the filtering, 150 traits entered downstream analysis (see **Supplementary Table 2** for all traits removed from analysis)

### LD score regression and further filtering

We applied LD score regression^3,47^ to estimate the heritability and genetic correlation across 150 traits. LD scores based on 1000 Genomes European data and the list of HapMap3 SNPs were downloaded from the LDSC GitHub repository. Only SNPs in HapMap3 were used to perform the regression.

To ensure the traits have a substantial genetic component, we remove 21 traits that do not have a heritability estimate that is significantly different from 0 (z statistic > 1.96). To further reduce the genetic overlaps of the traits, we remove 13 traits that have high genetic correlation (*r*_*g*_) with others. In short, if two traits have genetic correlation |*r*_*g*_| > 0.95, we remove the trait less enriched of genetic associations, quantified by the product of the sample size of the study (*N*) and the heritability of the trait (*h*^2^). The algorithm is as follows:

1. Sort the traits in decreasing order of *Nh*^2^.
2. Start from the trait with the highest *Nh*^2^, remove the traits that have |*r*_*g*_| > 0.95 with this trait.
3. Proceed to the trait with the next highest *Nh*^2^, repeat until no pairs of traits have |*r*_*g*_| > 0.95.

After the above filtering steps, we retain a final list of 116 traits for statistical analysis (**Supplementary Table 1**). We further restricted to the variants that are available for ≥ 50 traits, and in 1000 Genomes Phase 3 European sample with minor allele frequency (MAF) > 0.01. This leads to a total of 7,462,466 variants.

### Statistical Analysis Using fastASSET

The first step to study the patterns of pleiotropy is to quantify the level of pleiotropy across the genome. Previous studies often have quantified the pleiotropy of a SNP by counting the number of traits that reach *p* < 5 × 10^−8^ in individual trait analysis. This approach, however, is likely to miss many weaker associations. Here we described fastASSET, an accelerated version of the ASSET method which 1) detects SNPs associated with any trait in our collection (“global association”) and 2) quantifies the degree of pleiotropy of each SNP.

The original ASSET was built on top of meta-analysis across different phenotypes and search for the subset of traits with maximum meta-analysis z-statistic^7^. However, since we are analyzing a large number of traits, the orignal ASSET which searches through all subsets is computationally intractable. fastASSET reduces the computational burden by only searching among the traits with suggestive evidence of associations. We select the reduced list of traits by a liberal threshold and then adjust the z-statistics for the pre-selection to avoid “double-dipping”. The steps of the algorithm are as follows: 1) we first cluster the traits based on sample overlap and phenotypic correlation, quantified by LD score regression intercept; we then de-correlate the z-statistics within each cluster using Cholesky decomposition. 2) we apply a liberal threshold (*p* < 0.05) to the de-correlated z-statistics to choose the traits that are moderately associated with the SNP; 3) we make an adjustment to the de-correlated z-statistics to account for the pre-selection step based on conditional normal distribution 4) we then transform the adjusted z-statistics back to the correlated scale using the Cholesky decomposition in step 1; 5) we finally carry out a two-sided subset search, as implemented in the original ASSET algorithm, using the adjusted z-statistics for selected traits. The algorithm returns a p-value for global association that accounts not only for multiple-testing burden associated with the subset search but also for the pre-selection step used to reduce the number of traits included in all subset search. See **Supplementary Notes** for details. Simulation studies shows that fastASSET has well controlled type I error rate (**Supplementary Table 3**), and comparable power to detect global associations to metaUSAT and metaMANOVA^6^ under large sample size (e.g. *N* ≥ 50*k*, **Supplementary Figure 2**). The power loss under smaller sample size (N=20k) is due to loss of information from the pre-screening step which is designed to speed up computation. However, metaUSAT and metaMANOVA showed moderate type I error inflation when analyzing a large number of traits. In addition, the ability of fastASSET to select associated traits and quantify pleiotropy, which is not possessed by other methods, gives it unique advantage in multi-trait association analysis. See **Supplementary Notes** for the simulation settings.

### Analysis of data for 116 traits

Since different studies are relatively independent, LD score regression^3^ analysis did not reflect substantial sample overlap across different studies. The clusters, based on sample overlap and phenotypic correlation are generally small and restricted to traits within the same study (**Supplementary Figure 1**). For a small proportion of the SNPs, a large number of traits pass the pre-selection threshold (single-trait p-value < 0.05) which makes the subsequent subset search computationally intractable. We removed a total of 16,686 SNPs that were associated with more than 16 traits in one direction (either positive or negative). Under the global null hypothesis of no association, the probability of observing such pattern is small (1.13 × 10^−8^) and thus we consider these SNPs to be pleiotropic though they are not further analyzed by ASSET. In fact, the majority of these SNPs can be tagged by SNPs that are analyzed by ASSET through LD (r^2^>0.5) and hence removing them does not lead to significant loss of information, except 309 SNPs from 74 independent regions (r^2^<0.1, >500kb apart). We choose the SNP with the largest number of traits that pass p<0.05 threshold as the index SNP for each locus.

However, we show in validation and functional analysis that they are less pleiotropic than the highly pleiotropic SNPs we identified (**Supplementary Figure 11**), and hence do not have a major impact on the characterization of pleiotropy.

Among the SNPs we analyzed by ASSET, we further removed 2,620 SNPs for which one of the one-sided ASSET p-values is smaller than the p-values from standard meta-analysis of the selected traits, indicating convergence issues in the underlying p-value approximation method. For each of the remaining 7,443,160 SNPs, fastASSET reports a p-value for global association and a set of traits associated with the SNP. For each SNP, we use the number of its associated traits reported by fastASSET as the metric of pleiotropy. We consider SNPs with fastASSET p-value *p* < 5 × 10^−8^ as genome-wide significant. LD clumping leads to 10,628 independently associated SNPs with *r*^2^ < 0.1. We then group these SNPs into 2,293 independent loci of which the lead SNP (lowest p-value within the locus) are at least 500kb apart. As a metric of signal density within each locus, we count the number of independently associated SNPs within 100kb of the lead SNP.

We are especially interested in trait-specific SNPs (associated with only 1 trait as identified by fastASSET) and secondarily highly pleiotropic variants with associated with >15 traits.

### Validation in UK Biobank

We download the UK Biobank (UKB) GWAS summary statistics from Neale lab^31^. Among the 2,293 index SNPs of the independent loci identified by fastASSET, 6 SNPs are not included in the UK Biobank summary statistics. For each of those SNPs, we identify a proxy SNP in the UKB summary statistics as the one with the highest *r*^2^ within 100kb of the lead SNP. If *r*^2^ < 0.8 between the lead SNP and the proxy SNP, we exclude the locus from the validation study in UKB. We successfully found proxies for 5 index SNPs and one other (rs12203592) does not have a proxy SNP with *r*^2^ ≥ 0.8.

Among the 11,934 summary level datasets published by Neale lab (round 2 GWAS, August 1st, 2018), we removed the GWAS results for age, sex and 22 datasets that do not have a phenotype code. For continuous traits, we use the summary dataset for the inverse rank normalized trait (variable type continuous_irnt) and discard the dataset for the raw trait (variable type continuous_raw). We only keep the datasets for joint analyses across both sexes and discard the sex-stratified results. In addition, if there are multiple versions of GWASs for the same trait, we only keep the original version and remove the later versions. After filtering, we retain 4,114 summary level datasets for our validation study. For each the 2,292 index SNPs, we select the associated traits among the remaining 4,114 UKB traits by a per-SNP FDR threshold of 0.05.

### Relationship between pleiotropy and LD

To study the relationship between pleiotropy and LD, we estimate LD scores from 1000 Genomes Phase 3 EUR sample using SNPs with MAF>0.01 and 1 centiMorgan (cM) window. The calculation is performed using ldsc software^47^. Note that here we choose to estimate LD scores using all the SNPs in our analysis instead of using those downloaded from LDSC, which include only HapMap3 SNPs.

### eQTL status lookup and colocalization

To explore potential cis-regulatory mechanisms that may drive pleiotropy, we look up the lead SNPs of the 2,293 loci identified by fastASSET to explore the relationship between the pleiotropy and eQTL tissue/gene specificity. We accessed eQTL summary statistics for 49 tissues in GTEx v8^36^. We lift over the base pair coordinates from hg19 to hg38 to match the genome build of GTEx v8. For each of the lead variant, we count the number of tissues in which it is a significant eQTL for at least one gene with q-value < 0.05, and the total number of unique genes for which it is a significant eQTL regardless of tissues.

For each trait-specific locus, we conduct colocalization analysis between the GWAS signal for the corresponding trait and eQTL signals in GTEx v8. We restrict to protein-coding genes and gene-tissue pairs for which the lead SNP of the locus is a significant eQTL at q-value < 0.05. We use the SNPs within 50kb from the lead SNP as input to COLOC^42^. We consider the GWAS and eQTL signals to be colocalized if the posterior probability of colocalization (PP4) > 0.8.

### Chromatin state

To explore potential relationship between pleiotropy and chromatin states, we queried their chromatin state using 15-state chromHMM model^48^ in the Haplogreg v4.1 database^49^. The chromatin state was learned using a core set of 5 histone marks (H3K4me3, H3K4me1, H3K36me3, H3K27me3, H3K9me3) for 111 epigenomes from the Roadmap Epigenomics Project ^43^. We defined chromatin states with state number <=7 as an active and open chromatin state, including active transcription start site (1_TssA), flanking active TSS (2_ TssAFlnk), transcription at gene 5’ and 3’ (3_TxFlnk), strong transcription (4_Tx), weak transcription (5_TxWk), genic enhancers (6_EnhG) and enhancers (Enh). For each variant, we count the number of datasets in Roadmap whether it falls in active chromatin state as well as the number of broad categories provided by Roadmap (https://egg2.wustl.edu/roadmap/web_portal/meta.html). We also study and evaluate the association of pleiotropy with each active chromatin state separately. For example, we select the lead SNPs identified by fastASSET that are 1_TssA in at least one Roadmap dataset and investigate the relationship between the number of traits selected by fastASSET and the number of tissue/cell type groups in which this variant is in 1_TssA. We perform same analysis for other active chromatin states to learn the association. The tissue and cell type groups are defined by the Roadmap Epigenomics Consortium as provided in the link: (https://docs.google.com/spreadsheets/d/1yikGx4MsO9Ei36b64yOy9Vb6oPC5IBGlFbYEt-N6gOM/edit#gid=15).

In addition to chromHMM based model, we conduct similar analysis using the chromatin states learned by the IDEAS, an integrative and discriminative epigenome annotation system employing 2D genome segmentation method^32,33^ to jointly characterize the chromatin states across many different cell types. Since the IDEAS classification of chromatin states is marginally different from ChromHMM, we consider 10_TssA (Active Transcription Start Site), 8_TssAFlnk (Flanking Active TSS), 14_TssWk (Weak TSS), 5_Tx (Strong Transcription), 2_TxWk (Weak Transcription), 4_Enh (Enhancers), 6_EnhG (Genic Enhancers), 17_EnhGA (Active Genic Enhancers) as an active and open chromatin state. IDEAS algorithm here integrates epigenomes by preserving the position-dependency and cell type specific epigenetic events at fine scales^33^.

The IDEAS bigBed files were downloaded from http://bx.psu.edu/~yuzhang/Roadmap_ideas/ The bigBed files were converted to bed format using UCSC program bigBedToBed program fetched from the directory of binary utilities in http://hgdownload.cse.ucsc.edu/admin/exe/ The bed files thus processed were used for further downstream analysis, including the association of pleiotropy and epigenetic states. Blacklisted genomic region was filtered out where applicable as provided by ENCODE.

### Transcription factor binding sites

To understand potential relationship between pleiotropy transcription factor (TF) binding, we downloaded and referenced the JASPAR 2018^34^ and HOCOMOCO V11^35^, homosapiens comprehensive model collection of motif database. Intersection of variants with TF binding sites was performed by Bedtools v2.29.2^50^ to observe the association between SNPs and transcription factor binding sites.

To associate level of pleiotropy with TF binding profiles, the divide the 2,293 lead SNPs into two pleiotropy bins: 1) associated with 1-10 traits 2) associated with >10 traits. We calculate the proportion of SNPs in each bin that overlaps with transcription binding sites (TFBS). To explore whether this relationship is independent of the effects of other annotations, we fit the following logistic regression: (associated with >10 traits) ∼ (overlapping with TFBS) + (LD score) + (number of tissues for which the variant is an eQTL) + (number of eGenes) + (number of cell types for which the variant is in active chromatin state) and check the significance of the first predictor.

### Enhancer-gene connection

The activity-by-contact (ABC) model combines chromatin states with 3-dimensional contacts to map enhancers to their target genes ^51,52^. We download the enhancer-gene map for 131 human cell types and tissues constructed by the ABC model from the Engreitz lab website (https://www.engreitzlab.org/resources/). This dataset includes all enhancer-gene connections with ABC scores >= 0.015.

Among the lead SNPs that overlap with enhancers, we compute the correlation between the level of pleiotropy and the number of cell types for which the overlapping enhancer affects at least one target gene by the ABC model. We further compute the partial correlation adjusting for other annotations: LD score, number of tissues for which the variant is an eQTL, number of eGenes, number of cell types for which the variant is in active chromatin state.

### Matching trait-specific SNPs to highly pleiotropic SNPs

In the functional follow-up studies of 21 trait-specific SNPs, we match each of them to one SNP that is also associated with the given trait but show high-degree of pleiotropy (associated with >15 traits). For each trait-specific SNP (e.g. for Crohn’s disease), we examine the individual-trait p-value between the trait (e.g. Crohn’s disease) and all the highly pleiotropic SNPs and choose the one that shows the strongest association (lowest p-values) with the given trait (e.g Crohn’s disease). In this procedure, multiple trait-specific SNPs for one trait could be matched to the same highly pleiotropic SNP. This matching procedure applies to the examination of top 10 associated traits for lead SNPs (**Figure 4**), eQTL analysis (**Figure 5**) and chromatin state analysis (**Figure 6**).

## Code availability

fastASSET is available via the main ASSET R package (https://github.com/sbstatgen/ASSET, function *fast_asset*) and will be added to the Bioconductor package in the next release (https://bioconductor.org/packages/release/bioc/html/ASSET.html). The developmental version and other analysis scripts are available on https://github.com/gqi/fastASSET.

## Web resources

GRASP repository: https://grasp.nhlbi.nih.gov/Overview.aspx

LDSC: https://github.com/bulik/ldsc

GTEx Consortium: https://www.gtexportal.org/home/

Roadmap Epigenomics Project: https://egg2.wustl.edu/roadmap/web_portal/index.html

ASSET R package: https://bioconductor.org/packages/release/bioc/html/ASSET.html

Neale lab UK Biobank GWAS: http://www.nealelab.is/uk-biobank

JASPAR database for transcription factor binding profiles: http://jaspar.genereg.net/

ABC model: https://www.engreitzlab.org/resources/

## References

1. Buniello, A. et al. The NHGRI-EBI GWAS Catalog of published genome-wide association studies, targeted arrays and summary statistics 2019. Nucleic Acids Res. 47, D1005–D1012 (2019).

2. Visscher, P. M. & Yang, J. A plethora of pleiotropy across complex traits. Nat. Genet. 48, 707–708 (2016).

3. Bulik-Sullivan, B. et al. An atlas of genetic correlations across human diseases and traits. Nat. Genet. 47, 1236–1241 (2015).

4. Watanabe, K. et al. A global overview of pleiotropy and genetic architecture in complex traits. Nat. Genet. 51, 1339–1348 (2019).

5. Pickrell, J. K. et al. Detection and interpretation of shared genetic influences on 42 human traits. Nat. Genet. 48, 709–717 (2016).

6. Ray, D. & Boehnke, M. Methods for meta-analysis of multiple traits using GWAS summary statistics. Genet. Epidemiol. 42, 134–145 (2018).

7. Bhattacharjee, S. et al. A Subset-Based Approach Improves Power and Interpretation for the Combined Analysis of Genetic Association Studies of Heterogeneous Traits. Am. J. Hum. Genet. 90, 821–835 (2012).

8. O’Reilly, P. F. et al. MultiPhen: Joint Model of Multiple Phenotypes Can Increase Discovery in GWAS. PLOS ONE 7, e34861 (2012).

9. Qi, G. & Chatterjee, N. Heritability informed power optimization (HIPO) leads to enhanced detection of genetic associations across multiple traits. PLOS Genet. 14, e1007549 (2018).

10. Turley, P. et al. Multi-trait analysis of genome-wide association summary statistics using MTAG. Nat. Genet. 50, 229–237 (2018).

11. Ray, D. & Chatterjee, N. A powerful method for pleiotropic analysis under composite null hypothesis identifies novel shared loci between Type 2 Diabetes and Prostate Cancer. PLOS Genet. 16, e1009218 (2020).

12. Davey Smith, G. & Ebrahim, S. ‘Mendelian randomization’: can genetic epidemiology contribute to understanding environmental determinants of disease?*. Int. J. Epidemiol. 32, 1–22 (2003).

13. Davey Smith, G. & Hemani, G. Mendelian randomization: genetic anchors for causal inference in epidemiological studies. Hum. Mol. Genet. 23, R89–R98 (2014).

14. Zheng, J. et al. Recent Developments in Mendelian Randomization Studies. Curr. Epidemiol. Rep. 4, 330–345 (2017).

15. Qi, G. & Chatterjee, N. Mendelian randomization analysis using mixture models for robust and efficient estimation of causal effects. Nat. Commun. 10, 1941 (2019).

16. Verma, A. et al. PheWAS and Beyond: The Landscape of Associations with Medical Diagnoses and Clinical Measures across 38,662 Individuals from Geisinger. Am. J. Hum. Genet. 102, 592–608 (2018).

17. Diogo, D. et al. Phenome-wide association studies across large population cohorts support drug target validation. Nat. Commun. 9, 4285 (2018).

18. Verma, A. et al. Human-Disease Phenotype Map Derived from PheWAS across 38,682 Individuals. Am. J. Hum. Genet. 104, 55–64 (2019).

19. Shen, X. et al. A phenome-wide association and Mendelian Randomisation study of polygenic risk for depression in UK Biobank. Nat. Commun. 11, 2301 (2020).

20. Solovieff, N., Cotsapas, C., Lee, P. H., Purcell, S. M. & Smoller, J. W. Pleiotropy in complex traits: challenges and strategies. Nat. Rev. Genet. 14, 483–495 (2013).

21. Stearns, F. W. One Hundred Years of Pleiotropy: A Retrospective. Genetics 186, 767–773 (2010).

22. Sakaue, S. et al. A global atlas of genetic associations of 220 deep phenotypes. medRxiv 2020.10.23.20213652 (2020) doi:10.1101/2020.10.23.20213652.

23. Jordan, D. M., Verbanck, M. & Do, R. HOPS: a quantitative score reveals pervasive horizontal pleiotropy in human genetic variation is driven by extreme polygenicity of human traits and diseases. Genome Biol. 20, 222 (2019).

24. Chen, C.-Y. et al. Analysis across Taiwan Biobank, Biobank Japan and UK Biobank identifies hundreds of novel loci for 36 quantitative traits. medRxiv 2021.04.12.21255236 (2021) doi:10.1101/2021.04.12.21255236.

25. Boyle, E. A., Li, Y. I. & Pritchard, J. K. An Expanded View of Complex Traits: From Polygenic to Omnigenic. Cell 169, 1177–1186 (2017).

26. Liu, X., Li, Y. I. & Pritchard, J. K. Trans Effects on Gene Expression Can Drive Omnigenic Inheritance. Cell 177, 1022-1034.e6 (2019).

27. Li, Y. R. et al. Meta-analysis of shared genetic architecture across ten pediatric autoimmune diseases. Nat. Med. 21, 1018–1027 (2015).

28. Kar, S. P. et al. Genome-Wide Meta-Analyses of Breast, Ovarian, and Prostate Cancer Association Studies Identify Multiple New Susceptibility Loci Shared by at Least Two Cancer Types. Cancer Discov. 6, 1052–1067 (2016).

29. Hung, R. J. et al. Cross Cancer Genomic Investigation of Inflammation Pathway for Five Common Cancers: Lung, Ovary, Prostate, Breast, and Colorectal Cancer. JNCI J. Natl. Cancer Inst. 107, djv246 (2015).

30. Bycroft, C. et al. The UK Biobank resource with deep phenotyping and genomic data. Nature 562, 203–209 (2018).

31. Neale, B. M. Rapid GWAS of Thousands of Phenotypes for 337,000 Samples in the UK Biobank. http://www.nealelab.is/uk-biobank (2018).

32. Zhang, Y., An, L., Yue, F. & Hardison, R. C. Jointly characterizing epigenetic dynamics across multiple human cell types. Nucleic Acids Res. 44, 6721–6731 (2016).

33. Zhang, Y. & Hardison, R. C. Accurate and reproducible functional maps in 127 human cell types via 2D genome segmentation. Nucleic Acids Res. 45, 9823–9836 (2017).

34. Khan, A. et al. JASPAR 2018: update of the open-access database of transcription factor binding profiles and its web framework. Nucleic Acids Res. 46, D260–D266 (2018).

35. Kulakovskiy, I. V. et al. HOCOMOCO: towards a complete collection of transcription factor binding models for human and mouse via large-scale ChIP-Seq analysis. Nucleic Acids Res. 46, D252–D259 (2018).

36. GTEx Consortium. The GTEx Consortium atlas of genetic regulatory effects across human tissues. Science 369, 1318–1330 (2020).

37. Stelzer, G. et al. The GeneCards Suite: From Gene Data Mining to Disease Genome Sequence Analyses. Curr. Protoc. Bioinforma. 54, 1.30.1-1.30.33 (2016).

38. Mandelkow, E.-M. & Mandelkow, E. Tau in Alzheimer’s disease. Trends Cell Biol. 8, 425–427 (1998).

39. Butcher Matthew J., Gjurich Breanne N., Phillips Tracy, & Galkina Elena V. The IL-17A/IL-17RA Axis Plays a Proatherogenic Role via the Regulation of Aortic Myeloid Cell Recruitment. Circ. Res. 110, 675–687 (2012).

40. Kabashima, K. et al. Prostaglandin E 2 –EP4 signaling initiates skin immune responses by promoting migration and maturation of Langerhans cells. Nat. Med. 9, 744–749 (2003).

41. Singh, S. B. et al. Human IRGM regulates autophagy and cell-autonomous immunity functions through mitochondria. Nat. Cell Biol. 12, 1154–1165 (2010).

42. Giambartolomei, C. et al. Bayesian Test for Colocalisation between Pairs of Genetic Association Studies Using Summary Statistics. PLOS Genet. 10, e1004383 (2014).

43. Kundaje, A. et al. Integrative analysis of 111 reference human epigenomes. Nature 518, 317–330 (2015).

44. Cummings, J. L. & Cole, G. Alzheimer Disease. JAMA 287, 2335–2338 (2002).

45. Iqbal, K., Liu, F., Gong, C.-X. & Grundke-Iqbal, I. Tau in Alzheimer Disease and Related Tauopathies. Curr. Alzheimer Res. 7, 656–664 (2010).

46. Boyle, P. A., Buchman, A. S., Wilson, R. S., Leurgans, S. E. & Bennett, D. A. Association of Muscle Strength With the Risk of Alzheimer Disease and the Rate of Cognitive Decline in Community-Dwelling Older Persons. Arch. Neurol. 66, 1339–1344 (2009).

47. Bulik-Sullivan, B. K. et al. LD Score regression distinguishes confounding from polygenicity in genome-wide association studies. Nat. Genet. 47, 291–295 (2015).

48. Ernst, J. & Kellis, M. Chromatin-state discovery and genome annotation with ChromHMM. Nat. Protoc. 12, 2478–2492 (2017).

49. Ward, L. D. & Kellis, M. HaploReg v4: systematic mining of putative causal variants, cell types, regulators and target genes for human complex traits and disease. Nucleic Acids Res. 44, D877–D881 (2016).

50. Quinlan, A. R. & Hall, I. M. BEDTools: a flexible suite of utilities for comparing genomic features. Bioinformatics 26, 841 (2010).

51. Fulco, C. P. et al. Activity-by-contact model of enhancer–promoter regulation from thousands of CRISPR perturbations. Nat. Genet. 51, 1664–1669 (2019).

52. Nasser, J. et al. Genome-wide enhancer maps link risk variants to disease genes. Nature 593, 238–243 (2021).

