## Supplementary Notes for "Genome-Wide Large-Scale Multi-Trait Analysis Characterizes Global Patterns of Pleiotropy and Unique Trait-Specific Variants"

#### 1 Details of the data filtering steps

We collect 358 summary-level datasets that cover a wide range of phenotypes, including 338 datasets from NIH GRASP repository and 20 from large GWAS consortia. We apply the following filtering steps to the datasets:

1. Remove datasets for which important information is missing. Scenarios include: 1) The genotypes are collected by a special chip without genome-wide coverage (metabochip, immunochip, exome array, etc). 2) The data only provide p-values for associations, but do not provide the direction of SNP effect on the trait. 3) The association testing was performed using nonstandard approaches or stratified analysis based on covariates. 4) The README file is missing hence the meaning of each column is unclear.

2. Remove studies with small sample size. For continuous traits, we remove studies with sample size >5,000; for binary traits, we remove studies with <5,000 cases or <5,000 controls.

3. Remove studies where >25% individuals are of non-European ancestry.

4. Remove duplicated traits. Since the GRASP repository includes studies spanning over a decade, there are often multiple studies for the same trait (e.g., multiple versions of height and BMI GWASs from the GIANT Consortium). Among the studies for the same trait, we keep the study with the largest sample size and discard the rest of them.

#### 2 fastASSET

The original ASSET is built on top of meta-analysis across different phenotypes (Bhattacharjee et al, 2012). Let $(\beta_k, s_k),\ k = 1, 2, ..., K$ denote the estimates of regression parameter and its standard error for a given SNP and $K$ traits. Let $z_k = \beta_k/s_k$ be the corresponding z-statistics. The standard meta-analysis uses the following statistic to test for association

$$z_{meta} = \sum_{k=1}^{K} w_k z_k$$

where $w_k = \frac{1/s_k}{1/\sqrt{\sum_{k=1}^{K} 1/s_k^2}}$.

Since it is unknown which traits are truly associated with the SNP, including traits that are not associated can lead to loss of power. Therefore, the ASSET method conducts a search through all subsets of traits and and finds the subset that gives maximum meta-analysis z-statistic. Denote the meta-analysis z-statistic based on subset $S$ by $Z(S)$, which can be expressed as

$$Z(S) = \sum_{k \in S} w_k z_k$$

This maximum meta-analysis z-statistic is used as the new test statistic:

$$Z_{max-meta} = max_{S \in \boldsymbol{S}} |Z(S)|$$

The p-value is approximated using discrete local maxima.

This test statistic is powerful when the effect of the SNP on all the traits are in the same direction, which is unlikely in our analysis due to the large number of traits. Hence we adopt the two-side approach:

1. Conduct subset search within the positive and negative z statistics separately.
2. Compute the conditional p-values given the signs of the z-statistics, denoted by $\tilde{P}^{+}_{DLM}$ and $\tilde{P}^{-}_{DLM}$.

3. Combine the p-values using Fisher's method

$$Z^{(2)}_{max-meta} = -2[\tilde{P}^{+}_{DLM} + \tilde{P}^{-}_{DLM}].$$

The new statistic $Z^{(2)}_{max-meta}$ follows a chi-squared distribution.

#### 2.1 Accelerating ASSET by pre-screening - fastASSET

When analyzing a large number of phenotypes, the number of subsets grows exponentially ($2^K - 1$ subsets for $K$ traits), which makes a full subset search intractable. To reduce the computational burden, we pre-screen the traits with suggestive association with the given SNP using a liberal p-value threshold, denoted by $p_{scr}$. We only conduct subset search among the traits with p-value $< p_{scr}$. Denote the correponding z-statistic by $z_{scr} = \Phi^{-1}(1 - p_{thr}/2)$.

Before the subset search, the z-statistics $z_k$ need to be adjusted for the pre-screening to prevent type I error inflation. Under the null hypothesis that the SNP is not associated with any of the given traits, if none of the studies have overlapping samples, the $z_k$ would be independent across studies and can be adjusted independently. The adjusted p-value is

$$p'_k = P(|Z| > |z_k| \mid |Z| > z_{thr}) = \frac{P(|Z| > |z_k|, |Z| > z_{thr})}{P(|Z| > z_{thr})} = \frac{P(|Z| > |z_k|, |Z| > z_{thr})}{p_{thr}}$$

which is equal to $P(|Z| > |z_k|)/p_{thr}$ if $|z_k| > z_{thr}$ and 1 if $|z_k| \leq z_{thr}$. Hence the adjusted z-statistic is $z'_k = \Phi^{-1}(1 - p'_k/2)$.

However, the data used for our study have complex sample overlaps, hence the $z_k$ of overlapping studies are no longer independent. To apply the above adjustment, we need to first de-correlate the z-statistics. The correlation of z-statistics due to sample overlap can be estimated using bivariate LD score regression:

$$E[z_{1j} z_{2j} | l_j] = \frac{\sqrt{N_1 N_2}\rho_g}{M} l_j + \rho^{(z)}_{12}$$

where $\rho^{(z)}_{12}$ is the correlation of $z_{1j}$ and $z_{2j}$ under the null hypothesis, and $l_j$ is the LD score. Since sample overlap usually occurs between traits in the same study or consortium, the correlation matrix of z-statistics $\boldsymbol{\rho}^{(z)} = \{\rho^{(z)}_{kl}\},\ k, l =$

$1, 2, ..., K$ roughly follows a block diagonal structure. First we use hierarchical clustering to cluster the studies based on distance matrix $1 - |\boldsymbol{\rho}^{(z)}|$. We cut the hierarchical clustering tree at 0.8 so that $\rho_{kl}^{(z)} < 0.2$ if studies $k$ and $l$ are in different clusters. We de-correlate the z-statistics within each cluster and ignore the between-cluster correlations.

Within each cluster, we first order the studies by effective sample size from smallest to largest. For continuous traits, the effective sample size is the total sample size; for binary traits, the effective sample size is defined as $\frac{N_{case}N_{control}}{N_{case}+N_{control}}$. Denote by $\boldsymbol{\rho}^{(z,t)}$ the LD score regression intercept matrix of the traits in cluster $t$. Denote by vector $\boldsymbol{Z}^{(t)}$ the z-statistics of the traits in cluster $t$. The de-correlation goes as follows:

1. Apply Cholesky decomposition to $\boldsymbol{\rho}^{(z,t)} = U^T U$.
2. Transform the z-statistics by $\tilde{\boldsymbol{Z}}^{(t)} = (U^T)^{-1}\boldsymbol{Z}^{(t)}$. Select the traits with
$$|\tilde{z}_k^{(t)}| > z_{thr}$$
and adjust the z-statistics using the steps detailed above. Denote the converted z-statistics by $\tilde{\boldsymbol{Z}}_{adj}^{(t)}$
3. Convert the z-statistics back to the original scale by $\boldsymbol{Z}_{adj}^{(t)} = U^T \tilde{\boldsymbol{Z}}_{adj}^{(t)}$. Leave the standard errors unchanged
4. Combine the z-statistics for the selected traits in all clusters as the input for ASSET analysis.

### 3 Simulation studies

#### 3.1 Null simulations

We conduct simulation studies to evaluate the type I error control of fastASSET. We showed in our previous publications that simulations models based on individual genotypes implies a model based on summary statistics (Qi and Chatterjee, 2018; Qi and Chatterjee, 2021). In particular, a null model implies that the summary statistics follow a multivariate normal distribution centered at 0. Hence we simulate the summary statistics directly as follows:

$$\hat{\beta}_j \sim N(\mathbf{0}, \Sigma/N). \tag{1}$$

Here $\hat{\beta}_j$ is a vector of GWAS regression coefficients for a single SNP across multiple traits. We set sample size $N = 20,000$ for all the traits, and hence the GWAS standard error is $1/\sqrt{N}$. Here $\Sigma$ is the covariance matrix of z-statistics under the null, incorporating both sample overlap and phenotypic correlation. Element $(k, l)$ of this matrix is $\frac{\rho_{kl} N_{kl}}{N}$, where $\rho_{kl}$ is the correlation between traits $k$ and $l$ and $N_{kl}$ is the number of overlapping samples. To obtain a realistic covariance structure, we set $\Sigma$ as the LD score regression intercept matrix for the 116 traits in our study. We apply fastASSET to the simulated summary statistics. Since we did not simulate LD structure, it is not possible to use LD score regression to estimate the $\boldsymbol{\rho}^{(z)}$ matrices. Instead, we directly use $\Sigma$ in place of $\boldsymbol{\rho}^{(z)}$ and treat it as known. Here we conduct 20 million independent simulations ($j = 1, 2, ..., 20,000,000$) to evaluate the type I error rate.

#### 3.2 Non-null simulations for power evaluation

To evaluate the power of fastASSET, we simulate the summary statistics as a sum of two vectors of length 116:

$$\hat{\beta}_j = \beta_j + e_j, \quad j = 1, 2, ..., 300$$

where $\beta_j$ is the true effect and $e_j$ is the error. The error vector is simulated from $e_j \sim N(\mathbf{0}, \Sigma/N)$, the same distribution used in the null simulations. We still use the same sample size $N$ for all traits, but vary it to 20k, 50k, 100k and 200k across different simulations.

To simulate the vector of true effects $\beta_j$, we randomly select $K$ traits (denote the indices of selected traits by $\mathcal{K}$) that have true association with the SNP. We vary $K$ to 2, 5, 10 or 15. We simulate the $k$-the element of $\beta_j$ by $\beta_{jk} \sim N(0, 0.02^2)$ if $k \in \mathcal{K}$, and $\beta_{jk} = 0$ otherwise. The normal distribution allows heterogeneous effect size across traits. Note that we only simulate 300 SNPs, which is sufficient for the evaluation of power and estimation of pleiotropy.

For both type I error and power, we compare fastASSET to metaUSAT and metaMANOVA (Ray and Boehnke, 2018), two statistical methods for multi-trait association testing.

### 4 References


- Bhattacharjee, Samsiddhi, et al. "A subset-based approach improves power and interpretation for the combined analysis of genetic association studies of heterogeneous traits." The American Journal of Human Genetics 90.5 (2012): 821-835.

- Qi, Guanghao, and Nilanjan Chatterjee. "Heritability informed power optimization (HIPO) leads to enhanced detection of genetic associations across multiple traits." PLoS genetics 14.10 (2018): e1007549.

- Qi, Guanghao, and Nilanjan Chatterjee. "A comprehensive evaluation of methods for Mendelian randomization using realistic simulations and an analysis of 38 biomarkers for risk of type 2 diabetes." International Journal of Epidemiology (2021).

- Ray, Debashree, and Michael Boehnke. "Methods for meta-analysis of multiple traits using GWAS summary statistics." Genetic epidemiology 42.2 (2018): 134-145.
