## Supplemental Figures for "Genome-Wide Large-Scale Multi-Trait Analysis Characterizes Global Patterns of Pleiotropy and Unique Trait-Specific Variants"

### Supplementary Figures

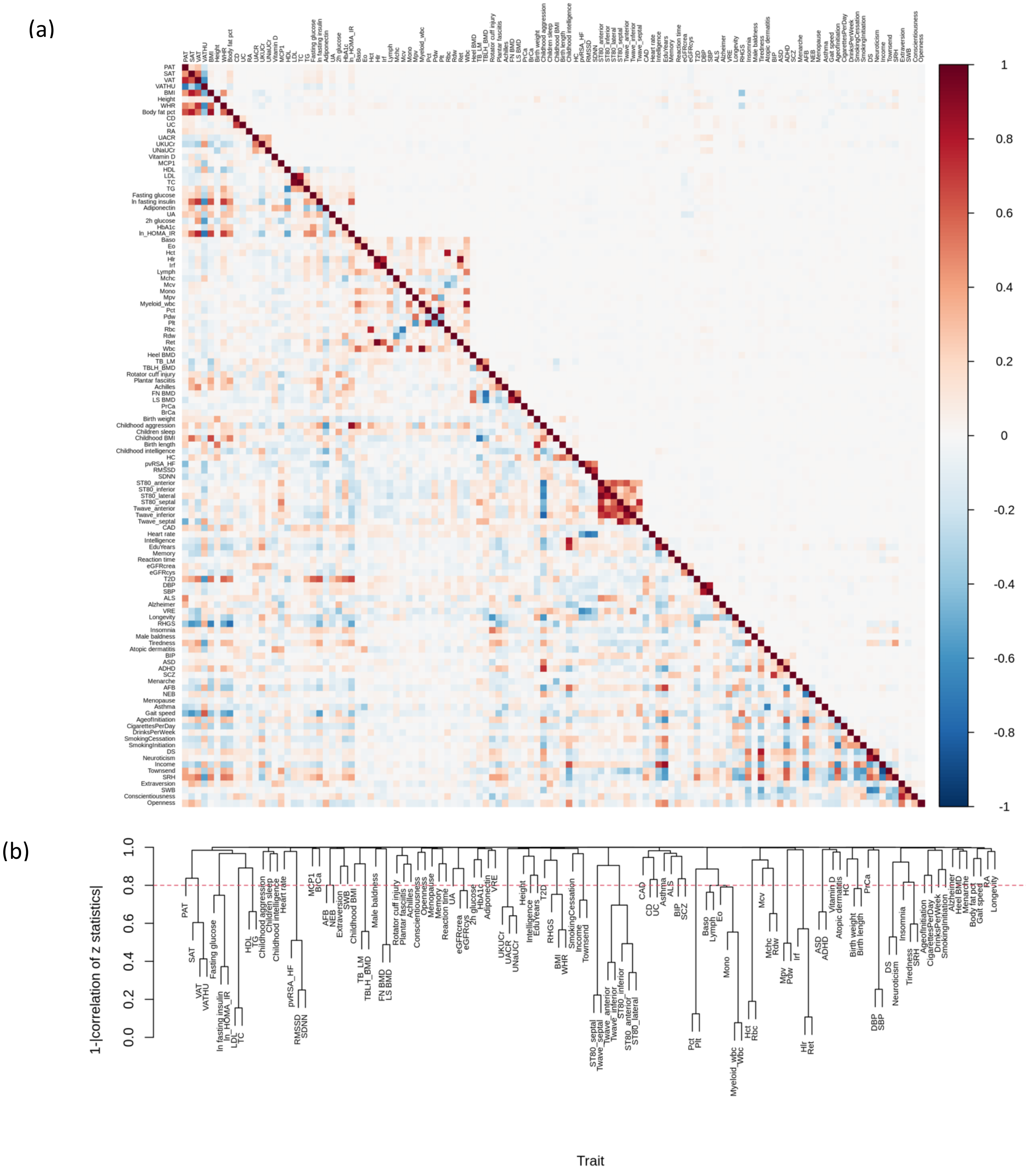


**Supplementary Figure 1.** (a) Genetic correlation (lower diagonal) across 116 traits estimated by LD score regression (LDSC) and correlation of z statistics under the global null hypothesis (upper diagonal, equal to LDSC intercept). Pairs of studies with substantially nonzero LDSC intercept needs to be de-correlated before ASSET analysis (Supplementary Notes). De-correlation is performed within clusters of traits defined by (b) hierarchical clustering based on LDSC intercept.

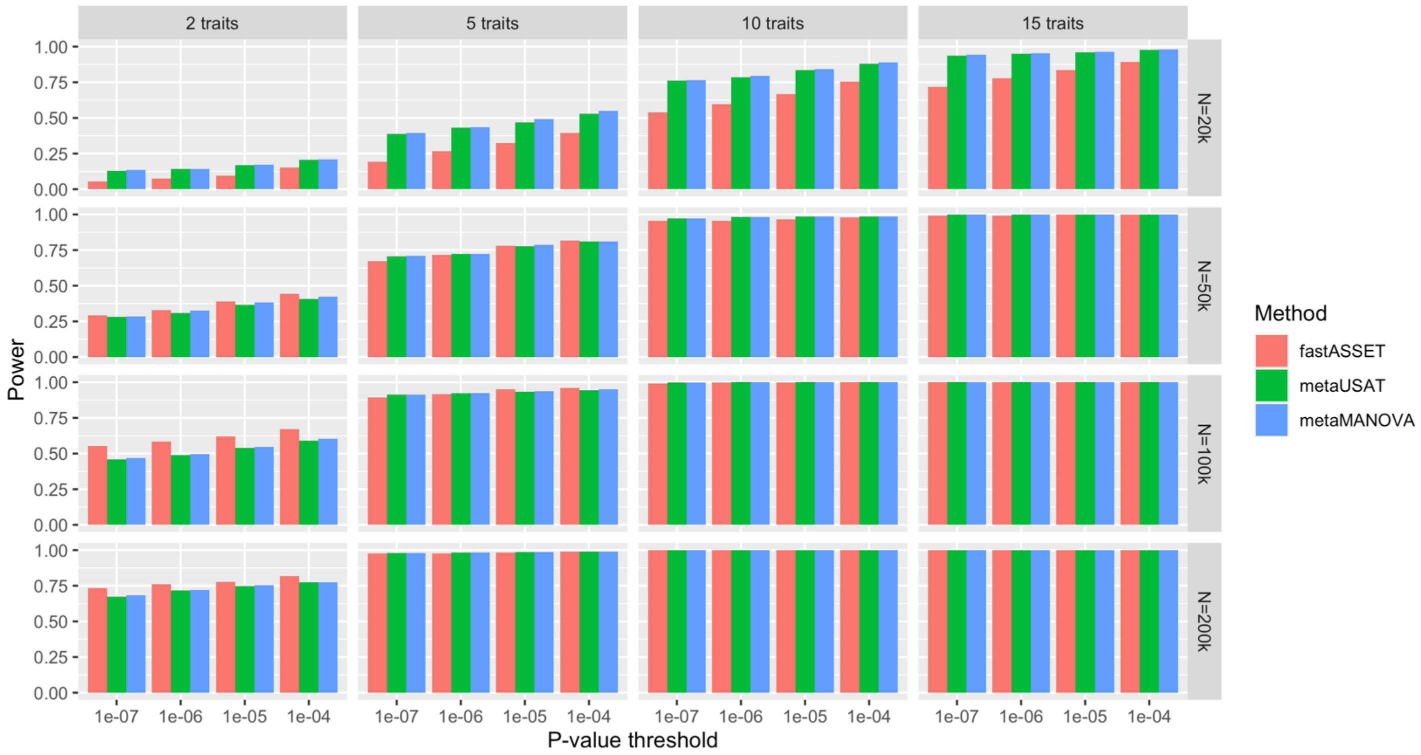


**Supplementary Figure 2. Power of fastASSET for detecting global association in simulation studies.** The null hypothesis is that the SNP is not associated with any of the trait under consideration. GWAS regression coefficients for one SNP $j$ and 116 traits ($\hat{\beta}_j$, vector of length 116) are simulated from model $\hat{\beta}_j = \beta_j + e_j$, where $e_j$ is the error term generated from multivariate normal distribution reflecting realistic correlation across traits. The term $\beta_j$ is simulated by first randomly selecting a set of 2, 5, 10 or 15 traits (columns) which have true associations with the SNP, followed by generating the effect size from $N(0, 0.02^2)$. The normal distribution allows heterogeneous effect size across traits. We vary the effect size N, assumed to be the same across all traits, to 20k, 50k, 100k and 200k (rows). This simulation procedure is repeated 300 times for each setting. The power of fastASSET is compared to that of metaUSAT and metaMANOVA. See Supplementary Notes for details of the simulation settings.

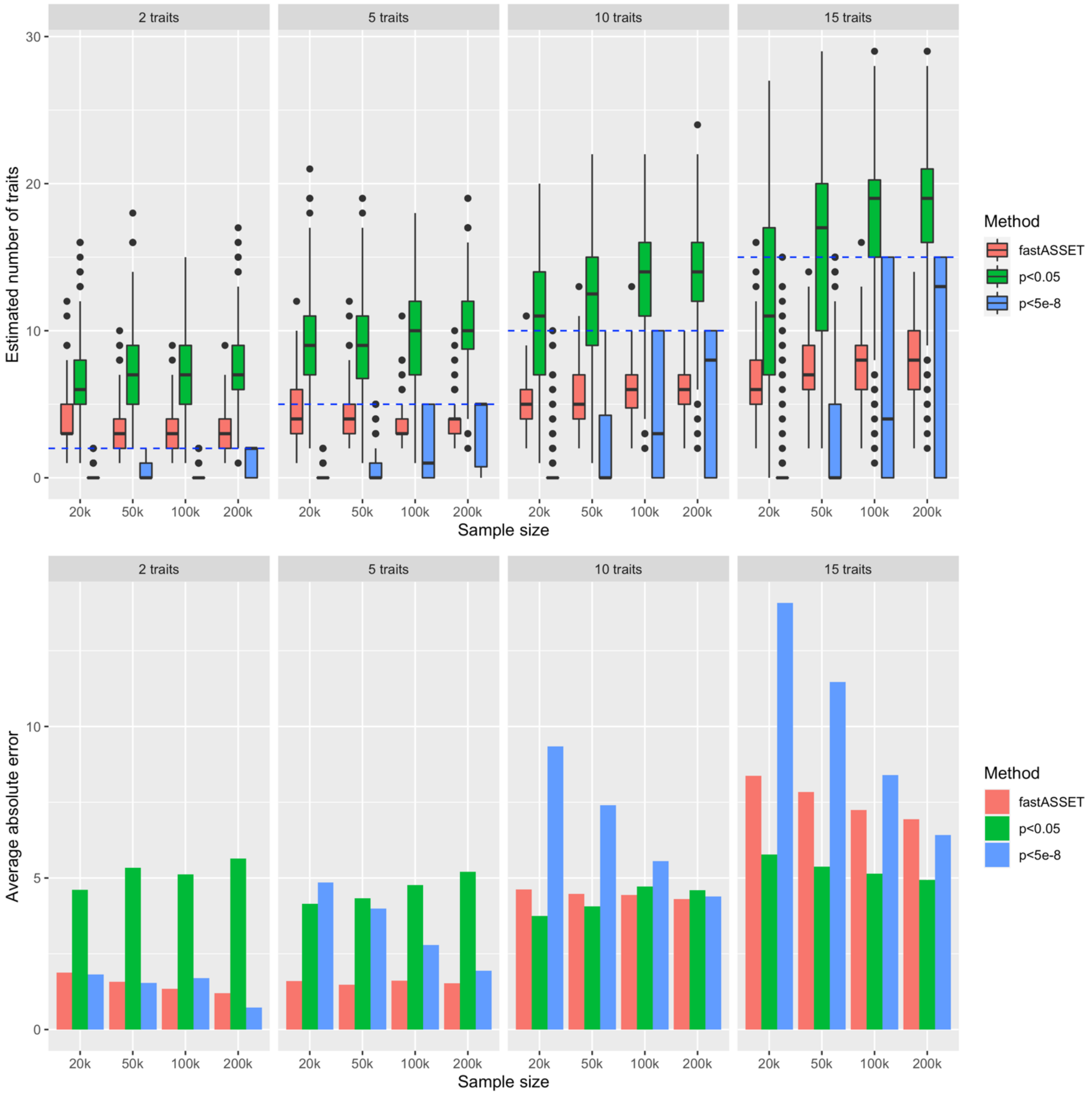


**Supplementary Figure 3. Accuracy of fastASSET for estimating of the level of pleiotropy in simulation studies.** Upper: estimated number of associated traits (level of pleiotropy) reported by fastASSET compared to counting the number of traits that reach p-value $< 0.05$ or $< 5 \times 10^{-8}$; title of each panel is the true number of associated traits, which is also marked by dashed blue lines. Lower: average absolute error for estimating level of pleiotropy, defined as average of |estimated number of associated traits – true number of associated traits| across 300 simulations. See legend of Supplementary Figure 2 or Supplementary Notes for description of the simulation settings.

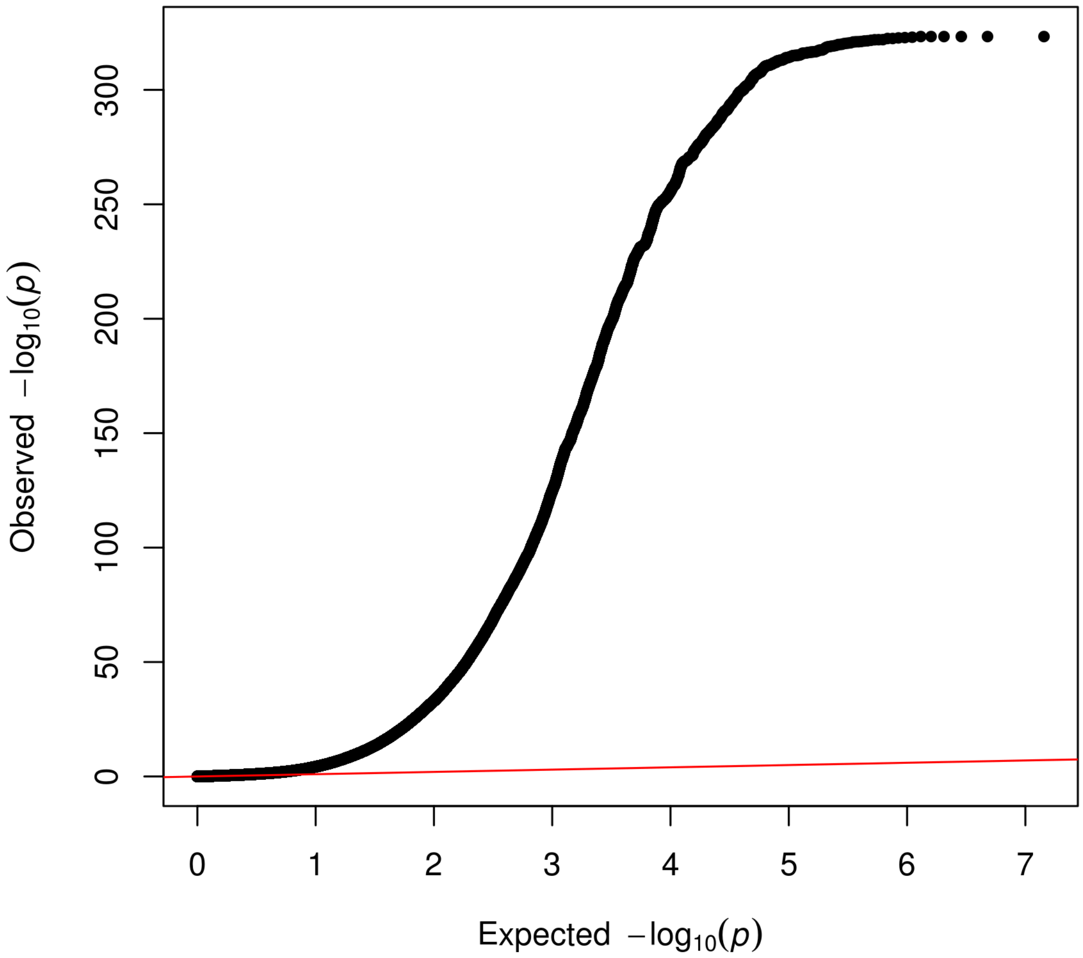


**Supplementary Figure 4. QQ plot for fastASSET p-values.**

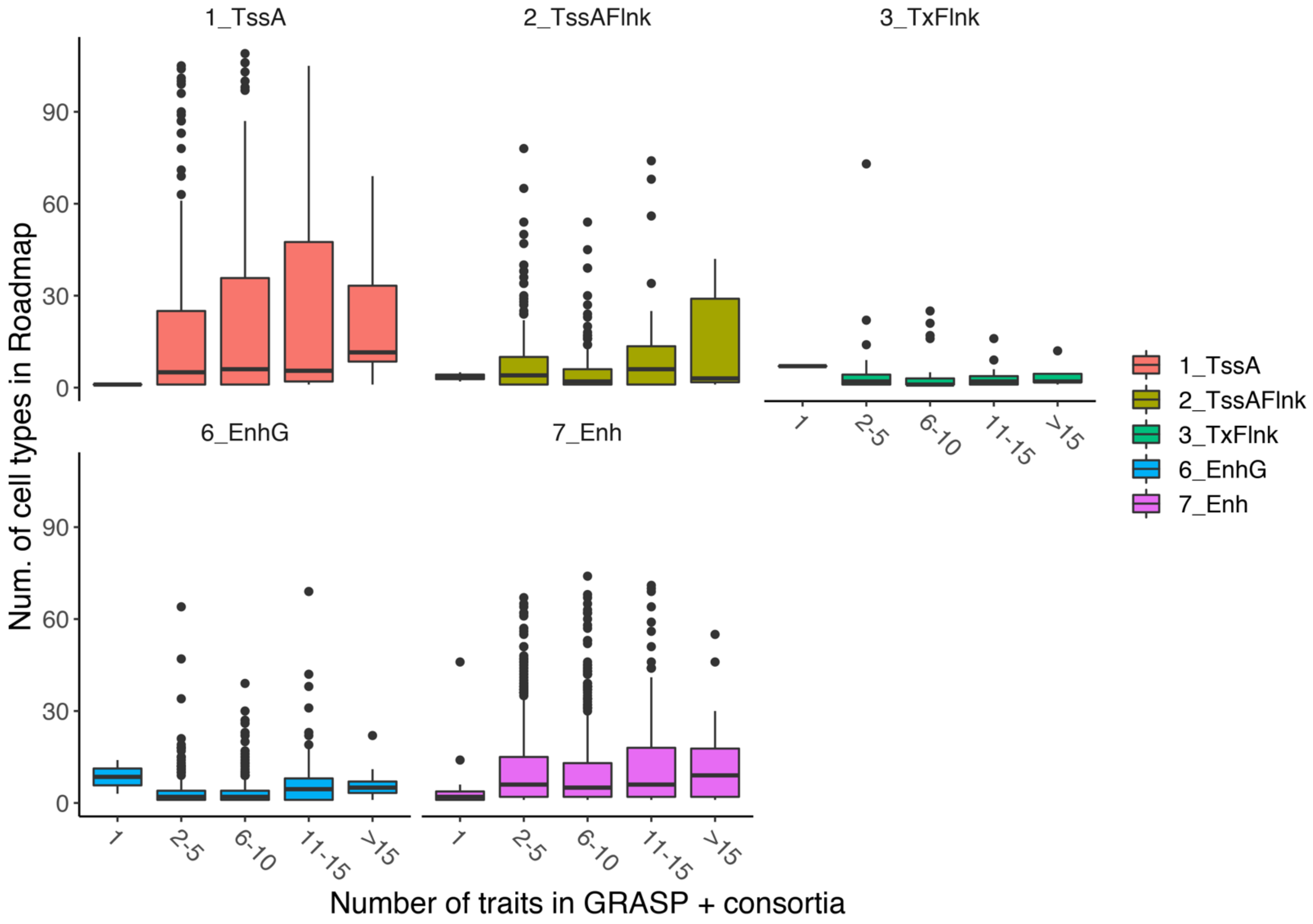


**Supplementary Figure 5. Relationship between pleiotropy and number of cell types where the SNP is in active chromatin state learned by ChromHMM.** Chromatin states are learned from Roadmap Epigenomics Consortium data.

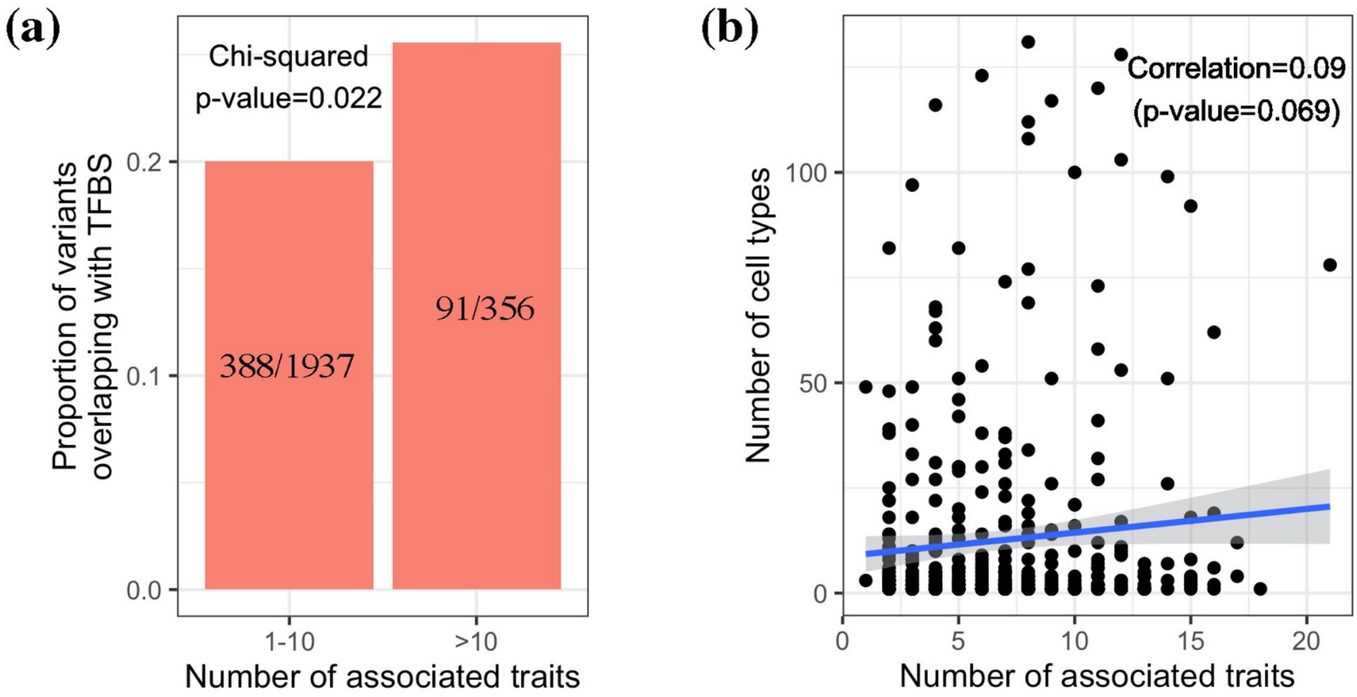


**Supplementary Figure 6: Relationship between pleiotropy and additional functional annotations.** (a) Overlap with transcription factor binding sites (TFBS) from the JASPAR and HOCOMOCO databases. (b) Cell type specificity of enhancer-gene connection identified by the activity-by-contact (ABC) model; the y axis is the number of cell types for which the enhancer overlapping with the SNP affects at least one target gene by the ABC model. After adjusting for other annotations (LD score, number of tissues for which the variant is an eQTL, number of eGenes, number of cell types for which the variant is in active chromatin state), the association in (a) remains significant (p-value=0.021), while the association in (b) disappears (partial correlation=0.02, p-value=0.65). See Methods for details.

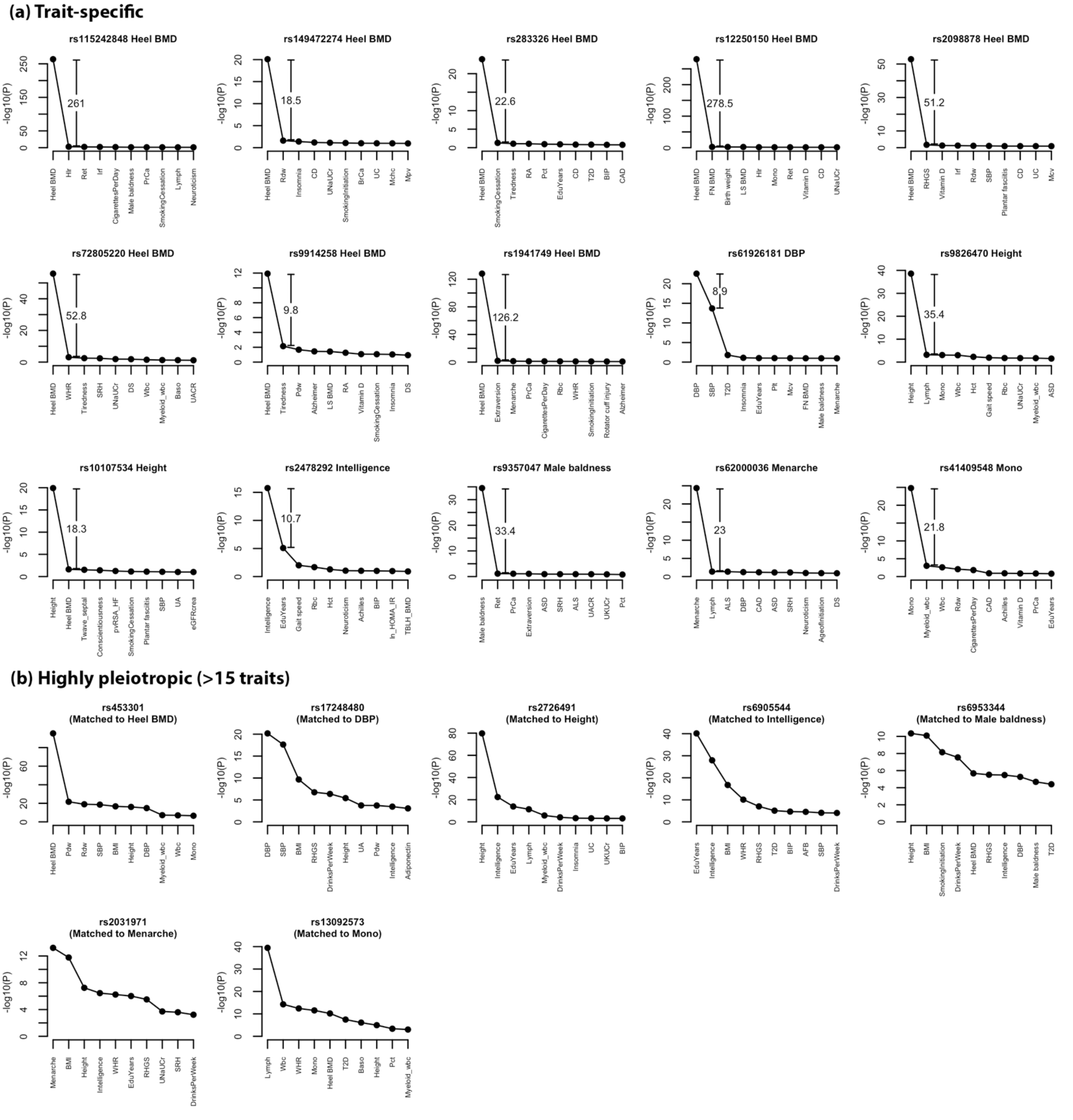


**Supplementary Figure 7. Top 10 associations of trait-specific SNPs for quantitative traits and matched highly pleiotropic SNPs.** Traits are ordered by descending order or -log10(p-value) in each panel. Each trait-specific SNP is matched to the highly pleiotropic SNP (>15 traits) that has the smallest association p-value with the trait (matched trait shown in the parentheses). See **Methods** for details of the matching procedure.

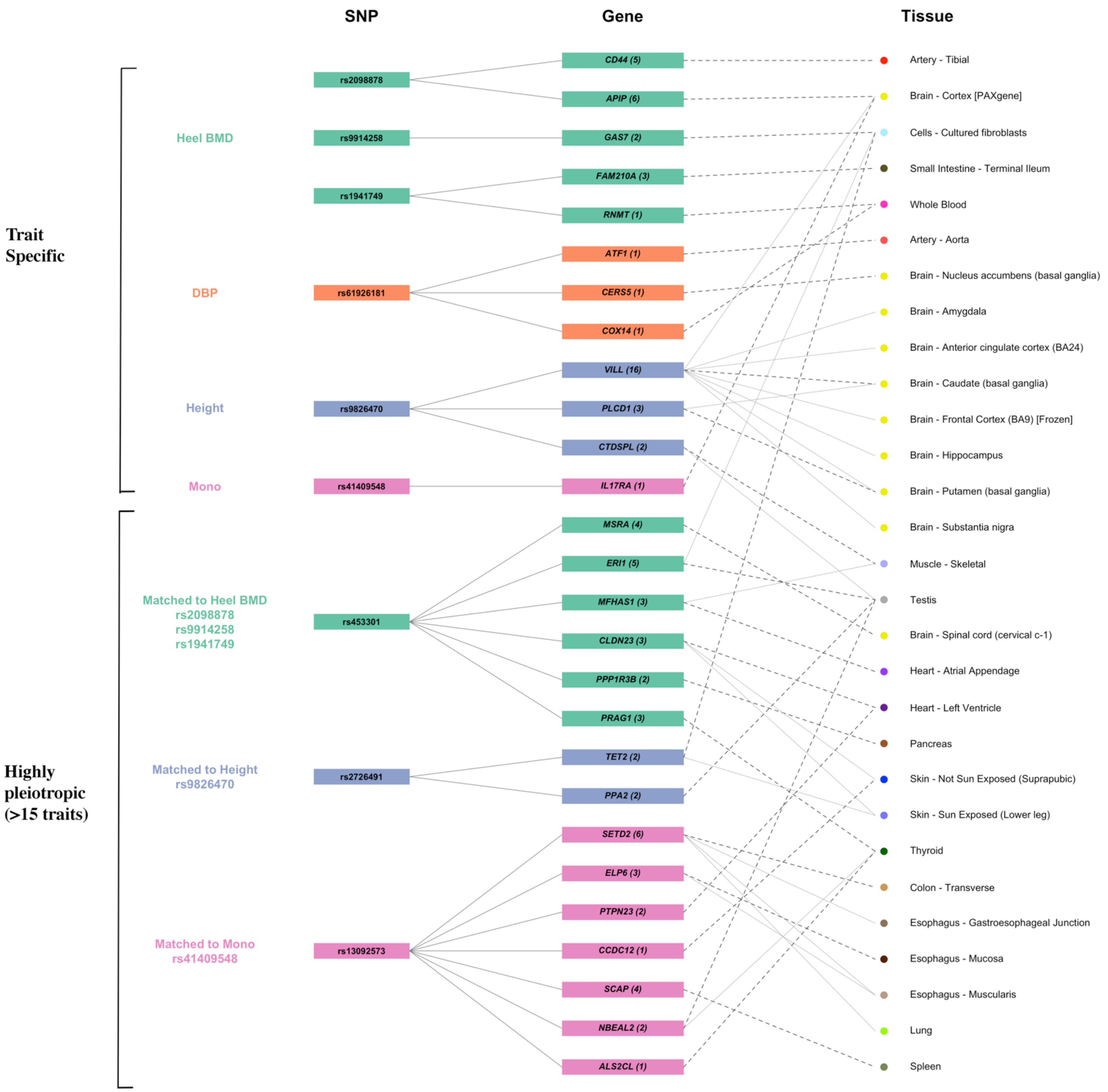


**Supplementary Figure 8. Cis-regulatory effects of trait-specific SNPs for quantitative traits and matched highly pleiotropic SNPs.** For each SNP, we show its protein-coding eGenes (q-value < 0.05) and the corresponding top tissues in GTEx v8. The top tissues for each variant-gene pair are defined as those with eQTL effect size >0.7*(the largest effect size among significant tissues for this variant-gene pair); the tissue harboring the largest effect is highlighted by darker dashed lines. In addition, we annotate each gene name by the total number of associated tissues (q-value < 0.05) regardless of effect size. Each trait-specific SNP is matched to the highly pleiotropic SNP (>15 traits) that have the strongest association with the trait (by p-value). See **Methods** for details of the matching procedure. The pleiotropic SNP matched to diastolic blood pressure (DBP) is not an eQTL for any gene-tissue pair and hence omitted. Pseudogenes and non-coding RNAs are excluded. BMD: bone mineral density; mono: monocyte count.

| Tissue | Samples | NES | p-value | m-value |
|---|---|---|---|---|
| Cells - EBV-transformed lymphocytes | 147 | 0.324 | 3.8e-4 | 0.818 |
| Artery - Aorta | 387 | 0.162 | 5.6e-7 | 1.00 |
| Spleen | 227 | 0.161 | 0.008 | 0.808 |
| Brain - Substantia nigra | 114 | 0.0912 | 0.3 | 0.470 |
| Lung | 515 | 0.0885 | 0.01 | 0.872 |
| Brain - Hippocampus | 165 | 0.0884 | 0.03 | 0.785 |
| Thyroid | 574 | 0.0740 | 0.04 | 0.881 |
| Whole Blood | 670 | 0.0630 | 1.0e-3 | 0.981 |
| Cells - Cultured fibroblasts | 483 | 0.0612 | 0.1 | 0.585 |
| Adrenal Gland | 233 | 0.0586 | 0.2 | 0.517 |
| Minor Salivary Gland | 144 | 0.0577 | 0.5 | 0.362 |
| Brain - Putamen (basal ganglia) | 170 | 0.0558 | 0.1 | 0.614 |
| Skin - Not Sun Exposed (Suprapubic) | 517 | 0.0558 | 0.02 | 0.765 |
| Artery - Tibial | 584 | 0.0538 | 0.05 | 0.701 |
| Brain - Spinal cord (cervical c-1) | 126 | 0.0514 | 0.5 | 0.323 |
| Uterus | 129 | 0.0468 | 0.6 | 0.323 |
| Small Intestine - Terminal Ileum | 174 | 0.0435 | 0.4 | 0.431 |
| Esophagus - Mucosa | 497 | 0.0381 | 0.2 | 0.459 |
| Artery - Coronary | 213 | 0.0379 | 0.4 | 0.378 |
| Brain - Nucleus accumbens (basal ganglia) | 202 | 0.0366 | 0.4 | 0.386 |
| Skin - Sun Exposed (Lower leg) | 605 | 0.0351 | 0.1 | 0.412 |
| Breast - Mammary Tissue | 396 | 0.0325 | 0.2 | 0.349 |
| Brain - Anterior cingulate cortex (BA24) | 147 | 0.0286 | 0.6 | 0.285 |
| Ovary | 167 | 0.0155 | 0.8 | 0.281 |
| Nerve - Tibial | 532 | 0.0147 | 0.5 | 0.0530 |
| Heart - Atrial Appendage | 372 | 0.0128 | 0.7 | 0.183 |
| Brain - Frontal Cortex (BA9) | 175 | 0.0120 | 0.8 | 0.233 |
| Brain - Caudate (basal ganglia) | 194 | 0.0104 | 0.8 | 0.179 |
| Testis | 322 | 0.00446 | 0.9 | 0.162 |
| Liver | 208 | 0.00412 | 0.9 | 0.192 |
| Prostate | 221 | -0.00374 | 0.9 | 0.157 |
| Brain - Hypothalamus | 170 | -0.0151 | 0.7 | 0.0960 |
| Adipose - Visceral (Omentum) | 469 | -0.0191 | 0.5 | 0.0210 |
| Adipose - Subcutaneous | 581 | -0.0197 | 0.4 | 0.00100 |
| Muscle - Skeletal | 706 | -0.0216 | 0.3 | 0.00700 |
| Heart - Left Ventricle | 386 | -0.0229 | 0.7 | 0.127 |
| Esophagus - Gastroesophageal Junction | 330 | -0.0352 | 0.4 | 0.100 |
| Brain - Amygdala | 129 | -0.0358 | 0.4 | 0.0490 |
| Pituitary | 237 | -0.0465 | 0.4 | 0.137 |
| Stomach | 324 | -0.0532 | 0.3 | 0.0730 |
| Vagina | 141 | -0.0571 | 0.2 | 0.0180 |
| Colon - Transverse | 368 | -0.0707 | 0.08 | 0.0200 |
| Brain - Cerebellar Hemisphere | 175 | -0.0729 | 0.3 | 0.111 |
| Kidney - Cortex | 73 | -0.0778 | 0.3 | 0.131 |
| Brain - Cortex | 205 | -0.0896 | 0.03 | 0.0220 |
| Colon - Sigmoid | 318 | -0.105 | 0.02 | 0.00500 |
| Pancreas | 305 | -0.139 | 0.01 | 0.0410 |
| Esophagus - Muscularis | 465 | -0.155 | 3.9e-4 | 0.00200 |
| Brain - Cerebellum | 209 | -0.224 | 1.4e-4 | 0.00400 |

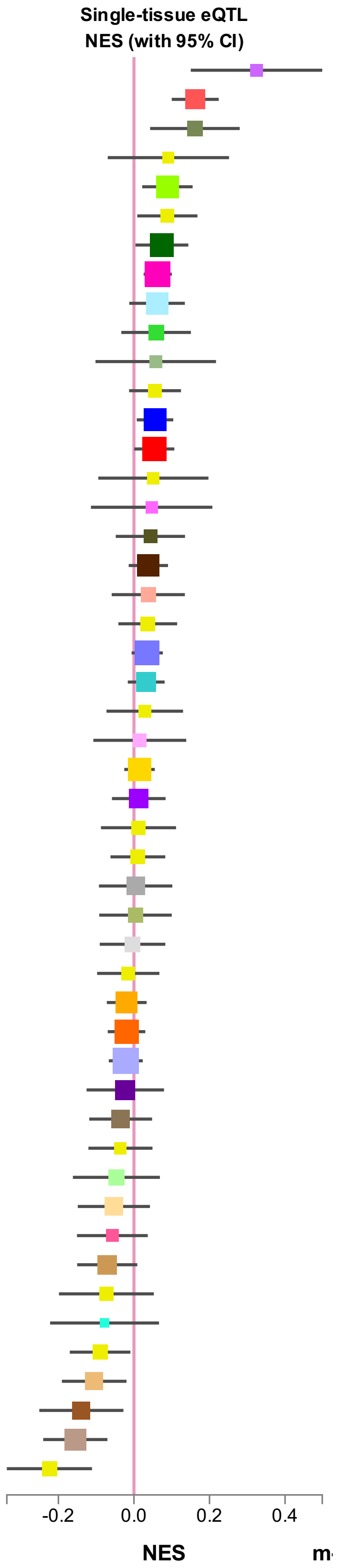


**Supplementary Figure 9. Multi-tissue eQTL plot for rs6733839 and BIN1 obtained from GTEx portal.** rs6733839 is a trait-specific SNP for Alzheimer’s disease.

| Tissue | Samples | NES | p-value | m-value |
|---|---|---|---|---|
| Liver | 208 | 0.0248 | 0.8 | 0.516 |
| Small Intestine - Terminal Ileum | 174 | 0.00804 | 1 | 0.738 |
| Kidney - Cortex | 73 | -0.00133 | 1 | 0.819 |
| Breast - Mammary Tissue | 396 | -0.00913 | 0.9 | 0.579 |
| Spleen | 227 | -0.0271 | 0.7 | 0.486 |
| Testis | 322 | -0.0289 | 0.8 | 0.806 |
| Cells - EBV-transformed lymphocytes | 147 | -0.0329 | 0.9 | 0.846 |
| Whole Blood | 670 | -0.0377 | 0.3 | 0.145 |
| Colon - Sigmoid | 318 | -0.0406 | 0.7 | 0.734 |
| Esophagus - Gastroesophageal Junction | 330 | -0.0666 | 0.5 | 0.769 |
| Stomach | 324 | -0.0840 | 0.4 | 0.875 |
| Artery - Tibial | 584 | -0.0893 | 0.4 | 0.879 |
| Adipose - Visceral (Omentum) | 469 | -0.0933 | 0.1 | 0.852 |
| Thyroid | 574 | -0.108 | 0.2 | 0.877 |
| Adipose - Subcutaneous | 581 | -0.122 | 0.1 | 0.961 |
| Heart - Atrial Appendage | 372 | -0.127 | 0.3 | 0.950 |
| Artery - Coronary | 213 | -0.133 | 0.4 | 0.926 |
| Esophagus - Muscularis | 465 | -0.138 | 0.1 | 0.962 |
| Lung | 515 | -0.142 | 0.06 | 0.967 |
| Brain - Putamen (basal ganglia) | 170 | -0.142 | 0.3 | 0.941 |
| Skin - Not Sun Exposed (Suprapubic) | 517 | -0.146 | 0.07 | 0.982 |
| Brain - Nucleus accumbens (basal ganglia) | 202 | -0.154 | 0.3 | 0.955 |
| Muscle - Skeletal | 706 | -0.164 | 0.01 | 1.00 |
| Skin - Sun Exposed (Lower leg) | 605 | -0.172 | 0.04 | 0.997 |
| Brain - Caudate (basal ganglia) | 194 | -0.176 | 0.2 | 0.964 |
| Colon - Transverse | 368 | -0.192 | 0.02 | 0.993 |
| Esophagus - Mucosa | 497 | -0.196 | 0.04 | 0.978 |
| Nerve - Tibial | 532 | -0.206 | 0.01 | 1.00 |
| Brain - Spinal cord (cervical c-1) | 126 | -0.225 | 0.4 | 0.929 |
| Prostate | 221 | -0.232 | 0.07 | 0.922 |
| Pancreas | 305 | -0.240 | 0.04 | 0.986 |
| Heart - Left Ventricle | 386 | -0.244 | 0.02 | 0.964 |
| Artery - Aorta | 387 | -0.264 | 0.01 | 1.00 |
| Brain - Anterior cingulate cortex (BA24) | 147 | -0.303 | 0.3 | 0.932 |
| Uterus | 129 | -0.329 | 0.3 | 0.950 |
| Brain - Hippocampus | 165 | -0.336 | 0.1 | 0.961 |
| Cells - Cultured fibroblasts | 483 | -0.345 | 0.01 | 0.990 |
| Brain - Amygdala | 129 | -0.351 | 0.2 | 0.946 |
| Minor Salivary Gland | 144 | -0.371 | 0.1 | 0.965 |
| Pituitary | 237 | -0.419 | 0.02 | 0.974 |
| Vagina | 141 | -0.445 | 0.09 | 0.983 |
| Adrenal Gland | 233 | -0.472 | 9.7e-4 | 0.996 |
| Ovary | 167 | -0.525 | 0.1 | 0.948 |
| Brain - Substantia nigra | 114 | -0.549 | 0.04 | 0.943 |
| Brain - Frontal Cortex (BA9) | 175 | -0.561 | 0.01 | 0.987 |
| Brain - Hypothalamus | 170 | -0.618 | 9.7e-4 | 0.991 |
| Brain - Cerebellar Hemisphere | 175 | -0.913 | 1.5e-3 | 0.997 |
| Brain - Cortex | 205 | -0.921 | 3.7e-6 | 0.997 |
| Brain - Cerebellum | 209 | -0.934 | 3.0e-4 | 0.998 |

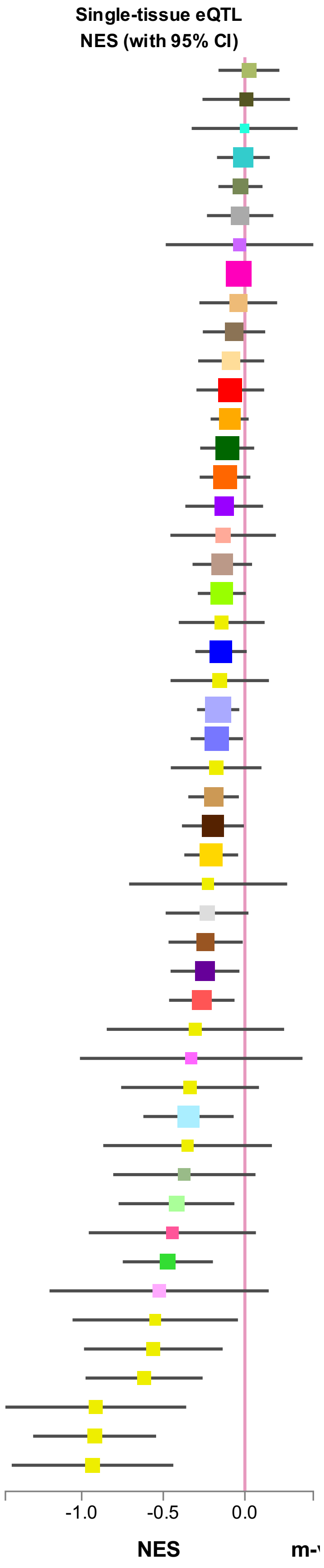


**Supplementary Figure 10. Multi-tissue eQTL plot for rs41409548 and IL17RA obtained from GTEx portal.** rs41409548 is a trait-specific SNP for monocyte count.

| Tissue | Samples | NES | p-value | m-value |
|---|---|---|---|---|
| Kidney - Cortex | 73 | 1.04 | 4.8e-5 | 1.00 |
| Artery - Tibial | 584 | 0.732 | 8.8e-18 | 1.00 |
| Adipose - Visceral (Omentum) | 469 | 0.690 | 5.8e-13 | 1.00 |
| Testis | 322 | 0.532 | 2.6e-5 | 1.00 |
| Breast - Mammary Tissue | 396 | 0.523 | 1.8e-7 | 1.00 |
| Ovary | 167 | 0.510 | 0.02 | 0.900 |
| Adipose - Subcutaneous | 581 | 0.450 | 3.9e-8 | 1.00 |
| Nerve - Tibial | 532 | 0.428 | 4.2e-6 | 1.00 |
| Uterus | 129 | 0.417 | 0.07 | 0.703 |
| Artery - Aorta | 387 | 0.401 | 1.0e-3 | 0.999 |
| Colon - Sigmoid | 318 | 0.236 | 0.05 | 0.185 |
| Artery - Coronary | 213 | 0.156 | 0.3 | 0.0450 |
| Prostate | 221 | 0.112 | 0.5 | 0.0300 |
| Thyroid | 574 | 0.0543 | 0.6 | 0.00 |
| Esophagus - Gastroesophageal Junction | 330 | 0.0226 | 0.9 | 0.00 |
| Adrenal Gland | 233 | - | - | - |
| Brain - Amygdala | 129 | - | - | - |
| Brain - Anterior cingulate cortex (BA24) | 147 | - | - | - |
| Brain - Caudate (basal ganglia) | 194 | - | - | - |
| Brain - Cerebellar Hemisphere | 175 | - | - | - |
| Brain - Cerebellum | 209 | - | - | - |
| Brain - Cortex | 205 | - | - | - |
| Brain - Frontal Cortex (BA9) | 175 | - | - | - |
| Brain - Hippocampus | 165 | - | - | - |
| Brain - Hypothalamus | 170 | - | - | - |
| Brain - Nucleus accumbens (basal ganglia) | 202 | - | - | - |
| Brain - Putamen (basal ganglia) | 170 | - | - | - |
| Brain - Spinal cord (cervical c-1) | 126 | - | - | - |
| Brain - Substantia nigra | 114 | - | - | - |
| Cells - Cultured fibroblasts | 483 | - | - | - |
| Heart - Atrial Appendage | 372 | - | - | - |
| Heart - Left Ventricle | 386 | - | - | - |
| Liver | 208 | - | - | - |
| Muscle - Skeletal | 706 | - | - | - |
| Pancreas | 305 | - | - | - |
| Pituitary | 237 | - | - | - |
| Skin - Not Sun Exposed (Suprapubic) | 517 | - | - | - |
| Skin - Sun Exposed (Lower leg) | 605 | - | - | - |
| Colon - Transverse | 368 | -0.0341 | 0.7 | 0.00 |
| Esophagus - Muscularis | 465 | -0.0456 | 0.6 | 0.00 |
| Cells - EBV-transformed lymphocytes | 147 | -0.0891 | 0.6 | 0.00800 |
| Lung | 515 | -0.109 | 0.3 | 0.00 |
| Stomach | 324 | -0.143 | 0.2 | 0.00 |
| Esophagus - Mucosa | 497 | -0.149 | 0.1 | 0.00 |
| Vagina | 141 | -0.187 | 0.4 | 0.00200 |
| Whole Blood | 670 | -0.211 | 2.3e-3 | 0.00 |
| Spleen | 227 | -0.396 | 0.006 | 0.00 |
| Minor Salivary Gland | 144 | -0.487 | 0.008 | 0.00 |
| Small Intestine - Terminal Ileum | 174 | -0.516 | 3.6e-5 | 0.00 |

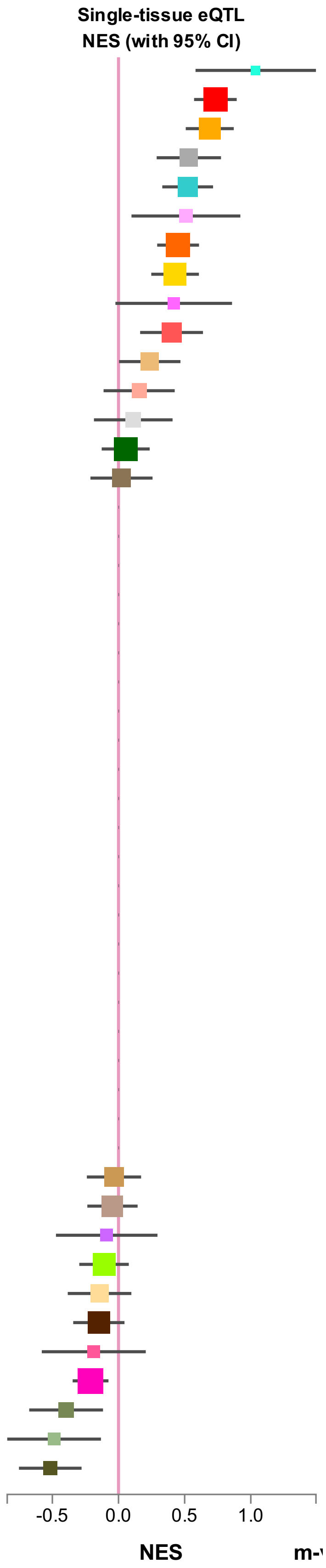


**Supplementary Figure 11. Multi-tissue eQTL plot for rs4958425 and IRGM obtained from GTEx portal.** rs4958425 is a trait-specific SNP for Crohn’s disease.

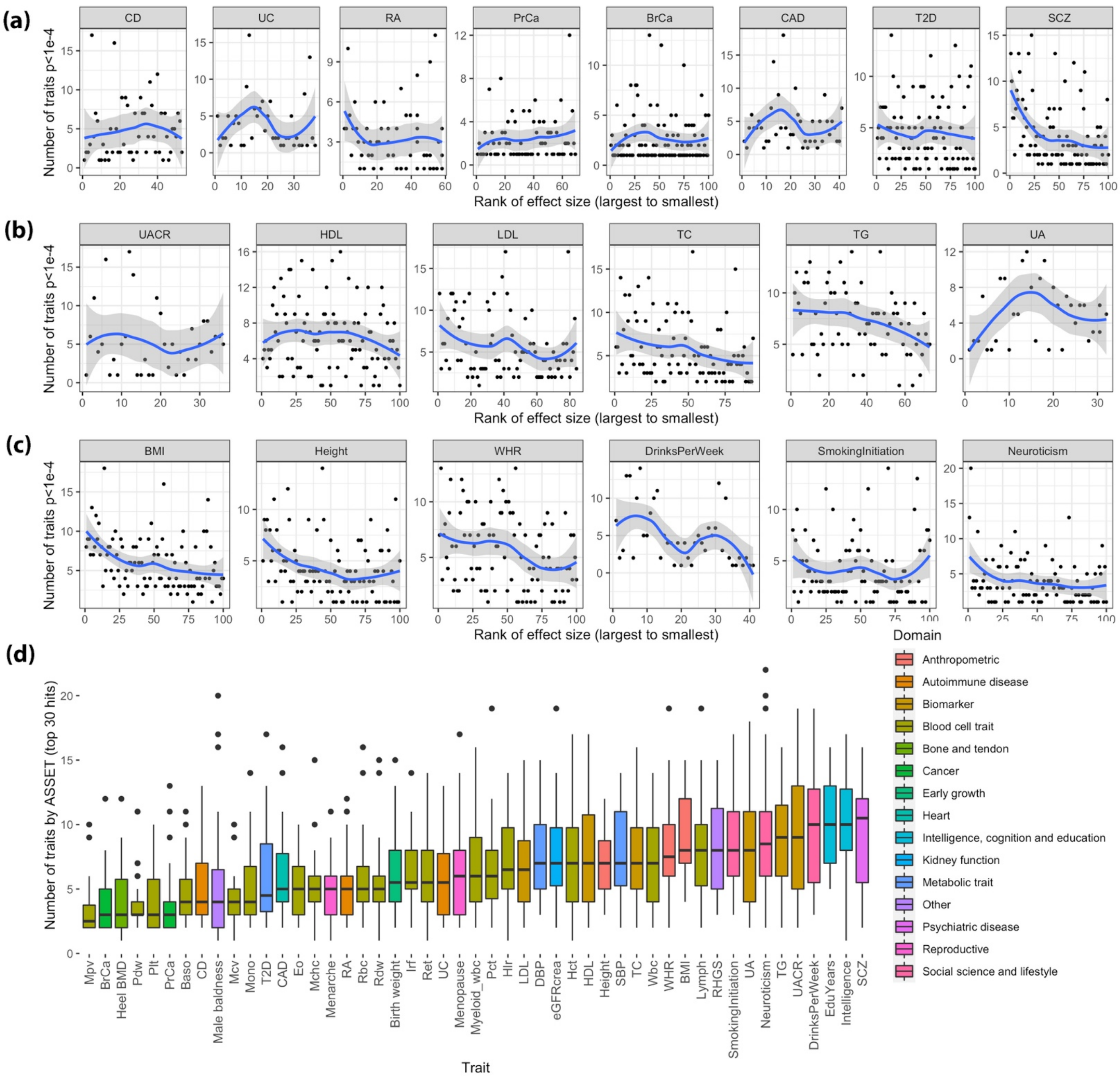


**Supplementary Figure 12. Patterns of pleiotropy and effect size across different traits.** (a)-(c) Relationship between level of pleiotropy and the effect size among lead SNP of loci associated with (a) complex diseases (b) blood and urine biomarkers with GWAS sample size >100,000 (c) anthropometric and social behavioral traits with sample size >100,000. Here the rank is based on the distribution of standardized effect size within the given trait. The standardized effect size is defined as $(\text{z statistic})/\sqrt{N}$, where N is the effective sample size (total sample size for quantitative traits; $\frac{N_{case}N_{control}}{N_{case}+N_{control}}$ for binary traits). In this analysis, for each trait we first identify those SNPs with $p < 5 \times 10^{-8}$ in the corresponding GWAS and perform LD clumping with $r^2 < 0.1$ and a genomic distance of >500kb. For each such index SNP that could be mapped to a trait of interest, we then identify all association with other traits based on a p-value cut-off of $p < 1 \times 10^{-4}$ in the underlying individual GWAS. The rank of the effect-sizes for the selected SNPs on the lead trait is then plotted against the number of other trait associations for these SNPs. The number of traits selected by fastASSET for top 30 lead SNPs ($p < 5 \times 10^{-8}$) associated with any given trait is also shown (d).

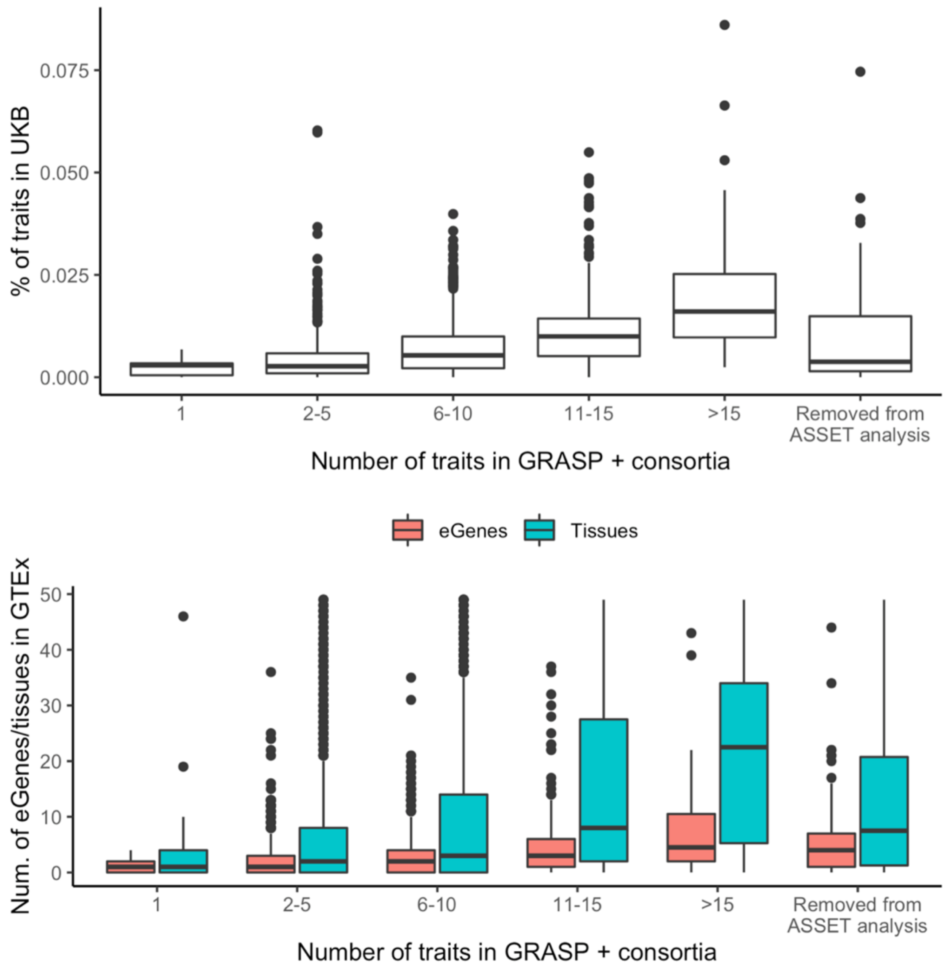


**Supplementary Figure 13. Patterns of pleiotropy for the SNPs that are removed from ASSET analysis compared to those in other categories of pleiotropy.** These SNPs represent 74 independent loci with $r^2 < 0.1$ and >500kb apart. Upper panel: levels of pleiotropy in UK Biobank; lower panel: number of tissues and genes for which a SNP is a significant eQTL in GTEx v8.
